# Convergent evolution of monocyte differentiation in adult skin instructs Langerhans cell identity

**DOI:** 10.1101/2023.11.13.566862

**Authors:** Anna Appios, James Davies, Sofia Sirvent, Stephen Henderson, Sébastien Trzebanski, Johannes Schroth, Morven L. Law, Inês Boal Carvalho, Howard Yuan-Hao Kan, Shreya Lovlekar, Christina Major, Andres Vallejo, Nigel J. Hall, Michael Ardern-Jones, Sian M. Henson, Elaine Emmerson, Steffen Jung, Marta E. Polak, Clare L. Bennett

## Abstract

Langerhans cells (LCs) maintain tissue and immunological homeostasis at the epidermal barrier site. They are unique among phagocytes in functioning both as embryo-derived, tissue-resident macrophages that influence skin innervation and repair, and as migrating professional antigen presenting cells, a capability classically assigned to dendritic cells (DCs). Here we report the mechanisms that determine this dual identity. Using ablation of embryo-derived LCs in murine adult skin and tracked differentiation of incoming monocyte-derived replacements, we reveal intrinsic intra-epidermal heterogeneity. We demonstrate that monocyte-dendritic cell progenitor (MDP)-derived monocytes are selected for survival in the skin environment. Within the epidermis, the hair follicle niche subsequently provides an initial site of LC commitment, likely via Notch signaling, prior to metabolic adaptation and survival of differentiated monocyte-derived LCs. In human skin, we show that embryo-derived (e)LCs in newborns retain transcriptional evidence of their macrophage origin, but this is superseded by distinct DC-like immune modules after post-natal expansion of eLCs. Thus, intrinsic and extrinsic adaptations to adult skin niches replicate conditioning of eLC at birth, permitting repair of the unique LC network.

## Introduction

Langerhans cells (LCs) are a unique and highly conserved population of mononuclear phagocytes that reside in the outer epidermis of the skin. Initially defined as prototypic dendritic cells (DCs) due to their potential to migrate to draining lymph nodes (LNs) and initiate T cell immunity (Jakob and Udey, 1999), subsequent fate mapping studies supported a common origin with tissue macrophages in other organs (Mass *et al*, 2016). As such, LCs are the only resident macrophage population that acquires the DC-like ability to migrate out of the tissue, and this dual nature is reflected in the breadth of LC function in the skin: as DC-like cells, LCs control T cell tolerance and function in LNs and the skin (Kaplan, 2017); by comparison, unlike DCs, LCs depend on CSF1R signalling for survival (Ginhoux *et al*., 2006), and there is increasing data to support more macrophage-like functions via interaction with peripheral nerves (Zhang *et al*., 2021) and promotion of angiogenesis during wound healing (Wasko *et al*., 2022). But the signals that control this functional dichotomy within the spatial context of intact skin remain poorly defined.

Tissue-resident macrophage (TRM) identity is largely imprinted by the local anatomical niche wherein instructive signals permit convergent differentiation and survival of long-lived, specialized resident cells irrespective of ontogeny (T’Jonck, Guilliams and Bonnardel, 2018; Blériot, Chakarov and Ginhoux, 2020). Environmental signals determine TRM identity via epigenetic regulation of specific transcription factor networks (Gosselin *et al*., 2014; Lavin *et al*., 2014), controlled by the transcription factor Zeb2 which is required for the expression of the unique gene programs that define identity and function within tissues (Scott *et al*., 2018). These niches have been carefully delineated in tissues such as the lungs and liver where interaction with local epithelia supports differentiation of resident macrophage populations (Bonnardel *et al*., 2019; McCowan *et al*., 2021). Implicit in these models is the concept of a single niche that provides all the essential requirements for TRM differentiation and survival as delineated by Guilliams *et al*: a physical scaffold; trophic factors to support maintenance of the network; and the signals to imprint a TRM identity unique to that site (Guilliams *et al*., 2020). But we questioned whether this model would also apply in the skin where monocytes must traverse the dermis upon leaving the blood and cross the epidermal basement membrane to populate the LC niche.

Murine fetal macrophage precursors expressing the quintessential protein CX3CR1, enter the early embryonic skin and differentiate into embryonic (e)LC-like cells within this environment (Hoeffel *et al*., 2012; Mass *et al*., 2016), but don’t mature into bona fide embryo-derived (e)LCs until after birth (Tripp *et al*., 2004; Chorro *et al*., 2009). By contrast, human eLCs differentiate within the epidermis before birth and histologically resemble adult cells by an estimated gestational age of 18 weeks (Meindl *et al*., 2009; Elbe-Bürger and Schuster, 2010). Entry of LC precursors into the BMP7- and TGFβ-rich environment of the epidermis results in activation of a Runx3- and Id2-dependent program of differentiation (Chopin *et al*., 2013; Mass *et al*., 2016; Kaplan, 2017; Strobl, Krump and Borek, 2019), defined by expression of the c-type lectin Langerin (CD207) and high expression of cell adhesion molecules including EpCAM and E-cadherin. Once resident, eLCs depend on IL34 for survival (Greter *et al*., 2012; Wang *et al*., 2012) and the network is maintained throughout life via local proliferation of mature LCs (Merad *et al*., 2002, 2004; Ghigo *et al*., 2013), independent of the adult circulation. In both mice and humans, eLC numbers are sparse at birth and the epidermis becomes fully populated following a proliferative burst within the first week of exposure to the external environment in mice (Chorro *et al*., 2009), and within 2 years of age in humans (Meindl *et al*., 2009), but we do not know if or how this post-natal transition shapes eLC identity and function.

Pathological destruction of the eLC network during graft-versus-host disease (GVHD) results in replacement of eLCs with donor bone marrow (BM)-derived LCs (Collin *et al*., 2006; Mielcarek *et al*., 2014). We and others have shown that acute inflammation and destruction of eLCs in murine models of GVHD or UV irradiation triggers influx of monocytes to the epidermis (Ginhoux *et al*., 2006; Seré *et al*., 2012; Santos e Sousa *et al*., 2018; Ferrer *et al*., 2019). By tracking monocyte differentiation, we demonstrated that epidermal monocytes undergo differentiation to EpCAM^+^CD207^neg^ precursors, which become long-lived EpCAM^+^CD207^+^ monocyte-derived (m)LCs that are transcriptionally indistinguishable from the cells they replace, are radio-resistant, and acquire DC-like functions of migration to LNs and priming of T cells (Ferrer *et al*., 2019). However, recent evidence suggests that heterogeneity within classical Ly6C^+^ monocytes results in some intrinsic bias in the cells that contribute to tissue-resident macrophage populations (Yáñez *et al*., 2017; Kwok *et al*., 2020). For instance, whilst all monocytes give rise to gut macrophages, in the lung, granulocyte-macrophage progenitor (GMP)-derived monocytes preferentially became Lyve1^+^MHCII^lo^ interstitial macrophages but monocyte-dendritic cell progenitor (MDP)-derived monocytes seeded the Lyve1^hi^MHCII^hi^ population (Trzebanski et al., 2023). These data suggest that monocyte ontogeny may shape tissue macrophage differentiation.

Utilizing our model of LC replacement, we have determined the process by which distinct cellular niches in the skin epidermis permit differentiation and survival of long-lived resident LCs. We demonstrate that a combination of BM monocyte ontogeny and environmental signals provided by the adult hair follicle niche instruct programs of LC development towards DC-like cells that replicate post-natal LC maturation in human skin. Together these data reveal mechanisms of convergent adaptation to the epidermal niche that imprints the unique LC identity in the skin.

## Results

### Single cell RNA-sequencing reveals monocyte-derived cell heterogeneity in the inflamed epidermis

To determine the molecular pathway(s) that resulted in successful tissue-residency and differentiation of mLCs in adult skin, we exploited our murine model of minor h-antigen mismatched HSCT, in which allogeneic T cells destroy resident eLCs (Merad *et al*. 2004; Santos e Sousa *et al*. 2018). In this model, mLCs replace the eLC network (Ferrer *et al*. 2019). We carried out single-cell RNA-sequencing (scRNA-seq) on sorted donor CD11b^+^MHCII^+^ cells isolated from the epidermis 3 weeks post-BMT with male antigen-specific Matahari (Mh) T cells (Valujskikh *et al*. 2002), using the 10X Genomics platform (Figure 1A, S1A). Analysis of the cells at this time point allowed us to map the spectrum of CD11b^high^ monocytes, CD11b^int^EpCAM^+^CD207^neg^ precursors and CD11b^int^EpCAM^+^CD207^+^ LCs we have previously defined in the epidermis (Ferrer *et al*. 2019). Dimensionality reduction and clustering of the cells demonstrated unexpected heterogeneity between donor BM-derived cells (Figure 1B). Several transcriptionally distinct clusters of cells surrounded a central collection of still distinct, but more convergent, clusters (Figure 1B). We used a parametric bootstrapping method sc-SHC (single-cell significance of hierarchical clustering) to demonstrate that the data, particularly to central clusters were not overfit and thus likely to be biologically relevant (Figure S2A)(Grabski, Street and Irizarry, 2023). Identification of the differentially expressed genes that defined the clusters (Figure 1C, S1B) revealed 3 populations of mLCs: resident (res) mLCs (*Cd207, Epcam, Mfge8*); cycling mLCs (*Top2a, Mki67*, but which also retained weakened expression of the res mLC signature); and migrating (mig) mLCs (*Cd83, Nr4a3, Ccr7*). Consistent with human LC data (Sirvent *et al*. 2020; Liu *et al*. 2021; Reynolds *et al*. 2021), mig mLC downregulated genes associated with LC identity (*Cd207, Epcam*) and instead expressed a generic monocyte-derived DC signature (Figure 1C, S1B) that showed the highest enrichment score for human migrating LCs across all clusters (Figure 1D) (Sirvent *et al*. 2020). These data suggest that some mLCs are constitutively primed for migration as observed in human steady state eLCs (Liu *et al*. 2021). We also identified a classical monocyte cluster (Mono; *Plac8, Ly6c2*) likely to have recently arrived in the epidermis, and two small populations of cells that shared monocyte/neutrophil activation markers, with signs of recent oxidative stress (S100a^+^ mono and Hmox1^+^ mono) (Figure 1C). The central overlapping clusters all shared macrophage identities with monocyte-derived cells found at other sites. The central cluster that shared most similarity to classical monocytes were defined as interferon-stimulatedgene monocytes (ISG mono; *Isg15, Ifit2, Ifit3*) as these genes were also evident in the classical monocyte cluster. Conversely, we observed a cluster of *Mrc1* (CD206)- and *Arg1*-expressing monocyte-derived macrophage-like cells (Mrc1^+^ mac) that resemble those found in the dermis (Figure 1C, S1B) (Kolter *et al*. 2019). These cells appeared to have differentiated along a default macrophage pathway to express canonical tissue macrophage genes. Intriguingly, we also detected a separate cluster of cells that were identified by their upregulation of *Ccl17, Mgl2, Dcstamp*, and *Itgax* genes linked to monocyte-derived DCs, and conventional type 2 DCs (cDC2s). (Figure 1C, S1B). Given the lack of clarity into the identity of these cells, we labelled them “converting monocyte-derived cells” (MC), in reference to a similar transitional population of cells in the pleural cavity recently described by the Allen lab (Finlay *et al*. 2023). Flow cytometry data validated the heterogeneous cell fates showing loss of monocytes and MC over time with expansion of mLCs and CD206^+^ macrophages (Figure 1E, F). Analysis of eLCs co-isolated from the GVHD skin at the 3 week time-point revealed 3 clusters of cells that resembled those identified in human skin (Liu *et al*. 2021): clusters 1 and 2 were defined as resident eLCs while the third cluster mirrored *Ccl22*^+^*Nr4a3*^+^ migrating mLCs (Figure S2B).

**Fig. 1.**
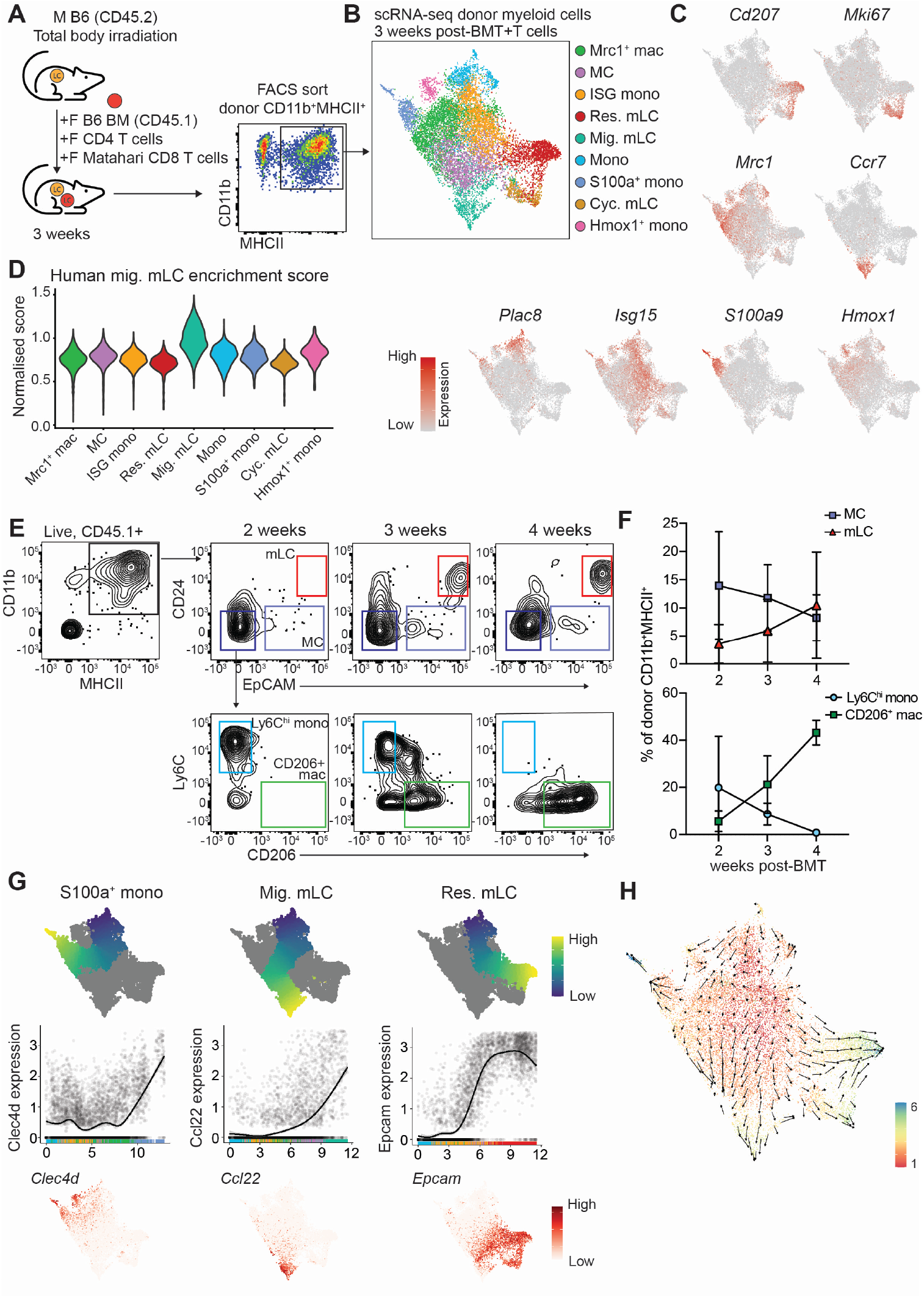
scRNA-seq reveals monocyte-derived cell heterogeneity in the GVHD epidermis. **A**. Experimental design showing murine bone marrow transplant model and populations sorted for scRNA-seq. M, male; F, female; BM, bone marrow. For full gating strategy see figure S1A. **B**. UMAP and clustering of donor CD11b^+^MHCII^+^ cells analyzed by 10X scRNA-seq. Mac, macrophage; MC, monocyte-derived cell; res. mLC, resident monocyte-derived Langerhans cell; mig. mLC, migratory mLC; mono, monocyte; cyc. mLC, cycling mLC. **C**. Heatmap overlays showing expression of indicated genes across dataset. **D**. Violin plot showing enrichment scores for a human mig. LC gene signature across clusters. **E**. Representative flow plots showing donor CD11b^+^MHCII^+^ cells from GVHD murine epidermis at the indicated time-points following BMT+T cells. Gated on live, singlets, CD45.1^+^ (donor) cells. **F**. Quantification of populations indicated in (E). Data are represented as mean±SD, (n=2 for 2 weeks, 8 for 3 weeks, 2 for 4 weeks). Data are pooled from three independent experiments. **G**. Differentiation trajectories calculated using Slingshot overlaid onto UMAP from (B) (above), normalised expression of indicated genes (y-axis) across pseudotime (x-axis) for the indicated trajectories (middle) and feature plots showing normalised expression of indicated genes overlaid onto UMAP from (B) (below). **H**. RNA velocity analysis applied to data from (B). Arrow directions indicate inferred cell trajectory.

To better understand which monocytes became mLCs, we inferred the trajectory of monocyte development towards finite cell states using Slingshot and RNA velocity (Figures 1G and H). These analyses revealed S100a^+^ monocytes as one of the endpoint cell states, illustrated by expression of *Clec4d* (Figure 1G). The Slingshot pathway passes mainly via the Mrc1^+^ macrophages whilst RNA velocity also suggested that some of this may be due to direct differentiation of incoming monocytes via the Hmox1^+^ monocyte cluster (Figure 1H). A separate pathway towards migratory mLC was inferred by high expression of *Ccl22* in the absence of *Cd207* (Figures 1C and G); this distinct route is likely driven by the down regulation of *Cd207* which is not expressed by migratory mLC (Reynolds *et al*. 2021). By comparison, upregulation of *Epcam* defined those monocytes destined to become resident mLCs (Figure 1G). But, whilst differentiation towards resident mLC populations was clearly defined as an endpoint in latent time (Figure 1H), arrows indicating direction of differentiation predicted by RNA velocity displayed a degree of uncertainty within Mrc1^+^ macs and ISG monocytes which was resolved once cells expressed *Epcam* (Figure 1G). This apparent lack of commitment was illustrated by the short latent time and relative increase in unspliced versus spliced transcripts within ISG monos and Mrc1^+^ macs compared to committed mLC populations (Figure S2C). Focusing on differentiation towards resident mLCs, progenitor marker gene analysis demonstrated loss of monocyte-specific genes (*Ly6c2, Plac8*) and acquisition of LC-defining genes (*Cd207, Epcam, Mfge8*), those associated with cell adhesion (*Cldn1*) and production of non-inflammatory lipid mediators (*Ptgs1, Ltc4s, Lpar3, Hpgds*) (Figure S2D).

Thus, specification of a mLC fate in the adult epidermis occurs *in situ* in the skin, but not all monocytes receive these signals and some undergo a default macrophage differentiation. Rather than discrete points of cell fate decisions, our combined analyses reveal a continuum of genes across the central clusters which converge to program mLC development in some cells.

### Monocyte ontogeny determines mLC repopulation

Heterogeneity in epidermal monocyte fate may be explained by either intrinsic bias within incoming Ly6C^high^ monocytes and/or extrinsic programming within a specified tissue niche. To test the first possibility, we investigated the epidermal monocyte population in more detail. Subsetting and reclustering of the classical monocyte cluster revealed 3 clusters of cells (Figure 2A): most cells fell into clusters 1 and 2 while a small number of cells were grouped in cluster 3 (Figure 2B). Enrichment for gene signatures for GMP-Mos and MDP-Mos (Trzebanski *et al*. 2023), and expression of the defining genes, *Cd177* and *Slamf7* suggested that cluster 3 represented GMP-derived monocytes (GMP-Mos), whereas clusters 1 and 2 were MDP-derived cells (MDP-Mos) (Figure 2C). This identification was supported by direct comparison between clusters 2 and 3; cluster 3 cells expressed higher levels of *Sell* (encoding CD62L) and classical monocyte genes (*Plac8, Ly6a2*), whereas cluster 2 expressed *C1q* genes and *H2-Aa*, associated with MDP-Mos (Figure 2D). To better understand the macrophage/DC potential within monocyte subsets we compared expression of a defined panel of genes (Figure 2E): cluster 3 (GMP-Mos) appeared more macrophage-like with higher expression of the glutathione reductase (*Gsr*) and *Cx3cr1*; clusters 1 and 2 were distinguished by increased, but differential, expression of *Id2, Mgl2*, and *Batf3* suggesting a closer relationship with DC-like cells (Figure 2E). Notably, *Mgl2* and *Ccl17*, markers of MDP-Mo progeny in the lung (Trzebanski *et al*. 2023) were expressed within MDP-Mo clusters (Figures 2E and S3A) and also within the MC cluster from our scRNA-seq dataset (Figure S1B).

**Fig. 2.**
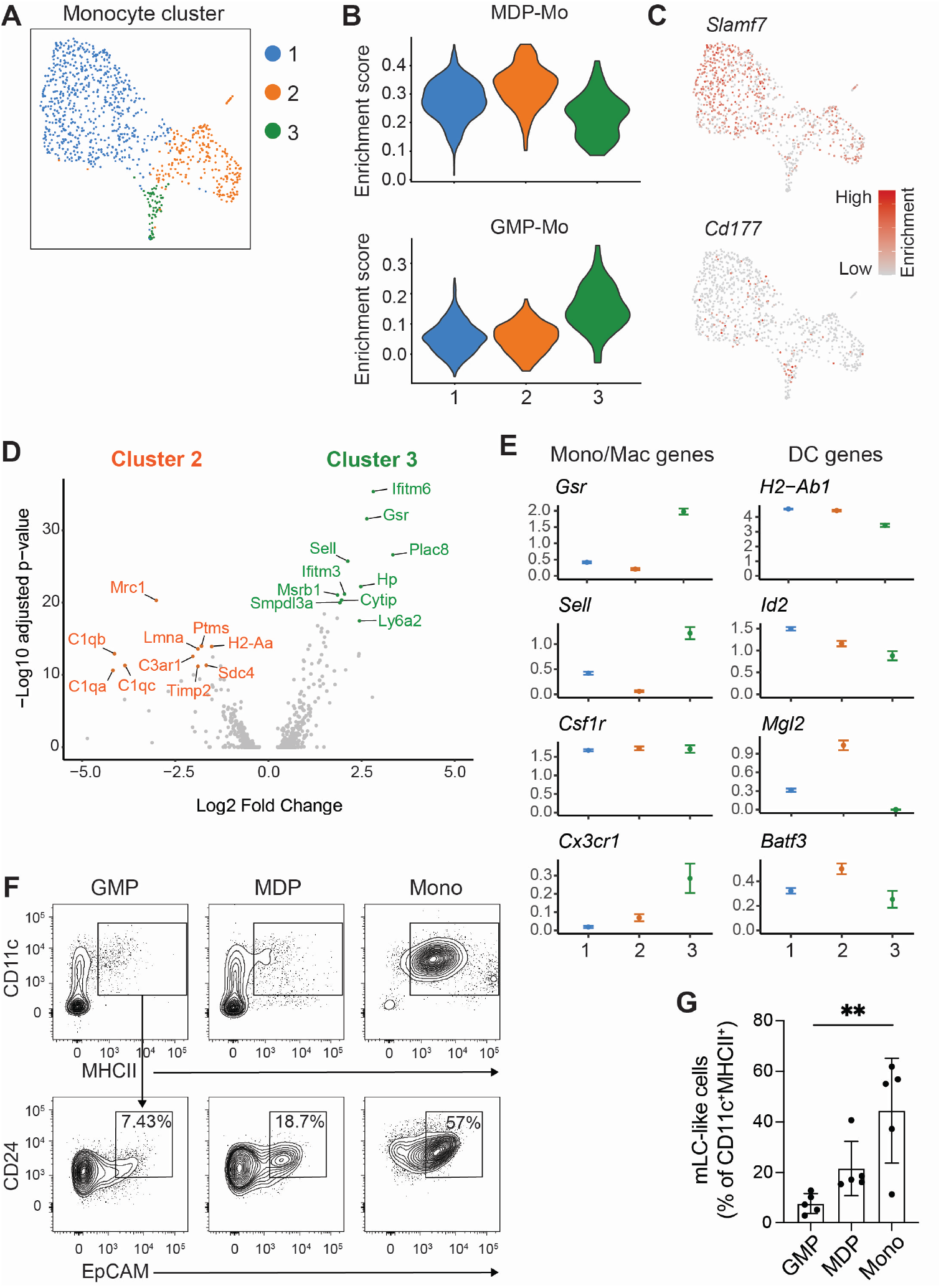
Intrinsically programmed DC-biased monocytes give rise to mLC. **A**. UMAP and sub-clustering of monocytes from GVHD epidermis. **B**. Violin plots showing enrichment scores for MDP-Mo (above) and GMP-Mo (below) gene signatures across clusters from (A). **C**. Heatmap overlays showing normalised expression of indicated genes. **D**. Volcano plot showing DEGs between cluster 2 and cluster 3 from (A). Top 10 significant DEGs are highlighted. **E**. Scatter plot of selected genes across monocyte clusters. **F** Representative flow plots showing GMP (left), MDP (middle) and monocytes (Mono, right) that were cultured for 6 days in the presence of GM-CSF, TGFβ and IL34. Gated on live, singlets, CD45^+^ cells. **G**. Bar graph showing the frequency of mLC-like cells (CD24^+^EpCAM^+^) as identified in (D). Data are represented as mean±SD (n=5 independent experiments). Statistical differences were assessed using a Friedman test, ^**^ p<0.01.

These data suggested that MDP-Mo were more likely to give rise to mLCs. Therefore, to test whether different monocytic precursors were intrinsically biased towards becoming mLC, we FACS-isolated GMP, MDP or total Ly6C^+^ monocytes from murine BM and cultured these cells with GM-CSF, TGFβ and IL34 to promote generation of CD24^+^EpCAM^+^ mLC-like cells (Figure S3B) (Chopin *et al*. 2013; Tabib *et al*. 2018; Ferrer *et al*. 2019). Figures 2F and G demonstrate that MDPs were more likely than GMPs to generate mLCs in culture, replicating differentiation of total Ly6C^+^ monocytes.

Together, these data highlight cell intrinsic differences between monocytes entering the epidermis; despite relative paucity in the blood (Trzebanski *et al*., 2023), MDP-Mo selectively expand within the epidermis with the potential to re-establish the nascent mLC network.

### Differentiating monocytes lose Zeb2-mediated macrophage identity to become mLCs but this does not require Ahr signaling

Analysis of the transcription factors that show the closest correlation with the Slingshot trajectory from monocytes to resident mLCs revealed that differentiation was dominated by loss of the tissue macrophage-determining factor *Zeb2* (Figure 3A). Expression of *Zeb2* was mutually exclusive to *Epcam*-expressing cells (Figures 3B, C), suggesting that, unlike other resident macrophage populations, a Zeb2-mediated macrophage fate is suppressed during specification of mLCs. Epidermal monocytes also downregulated expression of *Klf6*, linked to pro-inflammatory gene expression and therefore consistent with emergence of quiescent LCs (Goodman *et al*. 2016), as well as the transcription factors *Fos* and *Stat1*. Notably, *Zeb2* was highly negatively correlated with expression of the LC-defining transcription factors *Id2* and the aryl hydrocarbon receptor (*Ahr*) that are also upregulated by eLCs *in utero* (Mass *et al*. 2016) (Figures 3A-C). Mirroring DCs that migrate out of the skin, migratory mLCs were defined by upregulation of *Irf4, Rel* and *Nr4a3* (Figure S4A)(Baratin *et al*. 2015; Sirvent *et al*., 2020; Reynolds *et al*. 2021; Davies *et al*., 2022).

**Fig. 3.**
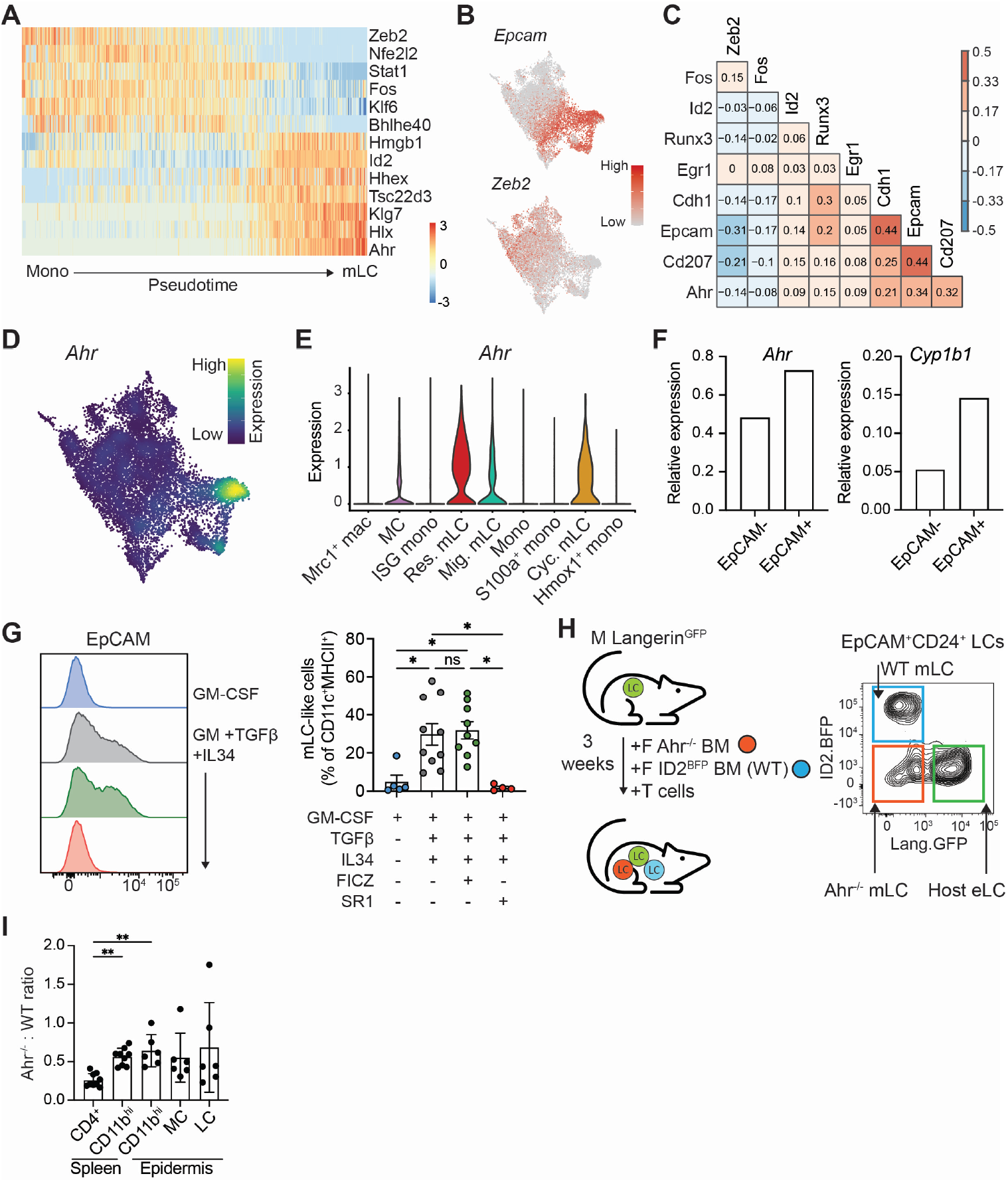
mLC differentiation is associated with loss of Zeb2 and upregulation of Ahr. **A**. Heatmap showing scaled gene expression of transcription factors that are differentially expressed along the differentiation trajectory (Pseudotime) from monocyte to res. mLC. **B**. Heatmap overlays showing normalised expression of indicated genes across UMAP from figure 1B. **C**. Correlation of selected LC-defining genes (y-axis) across all clusters of the scRNAseq dataset. **D**. Density plot showing expression of Ahr across cells from scRNA-seq dataset. **E**. Violin plot showing normalised expression of Ahr across clusters from scRNA-seq dataset. **F**. Relative expression of *Ahr* and *Cyp1b1* in CD11b^+^EpCAM^-^ and CD11b^+^EpCAM^+^ cells generated *in vitro* in the presence of FICZ. Expression is normalised to cells treated with GM-CSF+TGFβ+IL34 alone (n=2). **G**. Representative histogram (left) of EpCAM expression by monocytes cultured for 6 days under the indicated conditions and summary bar graph (right) of mLC-like cells generated from these conditions (for gating see Figure S4D). Data are represented as mean±SD (n=5 independent experiments). Statistical differences were assessed using Kruskal-Wallis test for multiple comparisons, ^*^ p<0.05. **H**. Experimental set up for competitive chimeras (left). Male Langerin^GFP^.B6 mice received a 1:1 mix of BM from female Ahr-replete (Ahr^+/+^.Id2^BFP^.B6 reporter mice, WT) or Ahrdeficient (Ahr^-/-^.B6) donors with Matahari T cells and donor chimerism was assessed in the epidermis and spleen 3 weeks post-transplant (right). M, male; F, female. **I**. Bar graph showing ratio of Ahr^-/-^ to WT frequencies of indicated cell types in spleen and epidermis of transplanted mice. Data are represented as mean±SD (n=6 for epidermis and 9 for spleen, from 2 or 3 independent experiments). Significant differences were assessed using a Kruskal-Wallis test, ^**^ p<0.01.

Previous work has shown that monocyte expression of Ahr biases differentiation towards moDCs rather than moMacs (Goudot *et al*. 2017). Ahr is not required for eLC development from pre-macrophages *in utero* (Figure S4B,C) (Esser, Rannug and Stockinger, 2009), but we questioned whether Ahr signaling could be important for MDP-Mo differentiation to mLCs in adult skin. Supporting this we also observed expression of Ahr by a small number of MCs, potentially indicating those cells en route to becoming mLCs (Figures 3D,E). To test whether Ahr signaling was required for monocyte differentiation *in vitro*, we generated mLCs in the presence or absence of the Ahr inhibitor SR1 or the agonist FICZ (Goudot *et al*. 2017). Both *Ahr* and *Cyp1b1* were augmented in EpCAM^+^ cells compared to EpCAM^neg^ cells in the presence of the agonist FICZ (Figure 3F), but activation of Ahr signaling did not result in an increase in the frequency of EpCAM^+^ mLC-like cells, probably due to Ahr ligands already present in culture media (Figures 3G and S4D) (Rannug and Fritsche, 2006). By contrast, inhibition of Ahr signaling ablated differentiation of EpCAM^+^ cells, demonstrating a requirement for mLC development *in vitro* (Figure 3G and S4D). Therefore, guided by these data we tested the requirement for Ahr *in vivo*. Exploiting the expression of ID2 and Langerin by LCs, we exploited Id2^BFP^ (Figure S4E,F)(Walker *et al*. 2019) and Langerin^GFP^ (Ferrer *et al*. 2019) reporter mice to generate competitive chimeras in which irradiated Langerin^GFP^.B6 males received a 1:1 mix of BM from female Ahr-replete (Ahr^+/+^.Id2^BFP^.B6 reporter mice) or Ahr-deficient (Ahr^-/-^.B6) donors with Matahari T cells (Figure 3H). Three weeks later the epidermis was analyzed for presence of mLCs and precursor populations. Consistent with the requirement for Ahr in CD4^+^ T cells (Esser, Rannug and Stockinger, 2009), Ahr-deficient BM cells did not contribute to repopulating splenic CD4^+^ T cells in chimeras (Figure 3I). We observed a slight bias towards Ahr-competent CD11b^+^ cells in the spleen across experiments (Figure 3I), however this ratio of Ahr-deficient to Ahr replete cells was maintained within our epidermal cells (Figure 3I). Therefore, these data demonstrated that Ahr signaling was not required for mLC differentiation *in vivo*.

Together these data suggest that loss of Zeb2 is a critical step for differentiation of mLCs. But, whilst Ahr signaling was required for monocyte differentiation *in vitro*, regulation by Ahr did not determine a mLC fate within adult skin.

### A unique follicular keratinocyte niche controls differentiation of mLCs

To understand the signals which regulated the transition from loss of a Zeb2-linked macrophage program to commitment to a LC identity, we next sought to define the mLC niche *in vivo*. Imaging of the skin after BMT with T cells revealed clustering of MHCII^+^ cells around hair follicles, an anatomical site previously associated with monocyte recruitment to the epidermis (Figure 4A)(Nagao *et al*. 2012). Therefore, to define where and how monocytes differentiated within potential epidermal niches we sorted CD45^neg^ keratinocytes (Figure S5) and CD11b^+^MHCII^+^ cells from the same epidermal samples 3 weeks post-BMT with T cells, performed scRNA-seq using the 10X platform and integrated the data with our existing CD11b^+^MHCII^+^ epidermal dataset. Clustering of CD45^neg^ cells followed by differential expression testing revealed a set of cluster-specific markers that corresponded with cluster markers of a previously published mouse scRNA-seq dataset (Joost *et al*., 2016). The comparison indicated a class of interfollicular epidermis-derived basal cells (Krt14^high^, Krt5^high^) and terminally-differentiated epidermal cells of the stratum spinosum (Krt10^high^), as well as a cluster that combined Krt79^high^ Krt17^high^ cells of the upper hair follicle with a small sub-cluster of Mgst1^+^ cells that were likely to come from the sebaceous gland (Figures 4B, C and S5B). Two other clusters were identified as cycling cells (*Mki67*) and putative *Cdc20*^+^ stem cells. To predict which keratinocytes could support differentiation of epidermal monocytes we analyzed expression of factors known to be required for monocyte or mLC survival (*Csf1, Il34, Bmp7*) and residency (*Tgfb1* and *2, Epcam*) (Kaplan, 2017). Figure 4D shows that while *Csf1* was not expressed by epidermal keratinocytes, *Il34*, which also binds the Csf1 receptor, was localised to Krt10^+^ terminally-differentiated cells, consistent with its role as a survival factor for the mature LC network within the interfollicular epidermis. We could only detect low levels of *Tgfb1* and *2* and *Bmp7* transcripts which may be due to lack of activation of TGFβ at this time point in GVHD skin but may also reflect poor detection of these genes at 10X Genomics sequencing depths. By contrast, *Epcam* was specifically expressed by *Krt79*^high^*Krt17*^high^ upper hair follicle cells. Expression of the cell adhesion molecule EpCAM, is associated with residency of monocytes within the alveolar space and differentiation to alveolar macrophages (McCowan *et al*. 2021), and EpCAM expression demarcates isthmus region epithelial cells of the upper hair follicle which express CCL2 (Nagao *et al*. 2012). Therefore, we postulated that this localised area may provide a niche for differentiating EpCAM^+^ LC precursors. Protein analysis demonstrated high levels of EpCAM on follicular epithelium, which was transiently decreased at the peak of T cell-mediated pathology in this model (Figure 4E,F) (Santos e Sousa *et al*. 2018; Ferrer *et al*. 2019), suggesting that adhesion to this niche could contribute to the bottleneck in mLC differentiation that we observed in our previous study (Ferrer *et al*. 2019). Using the LIANA software (Dimitrov *et al*. 2022), we predicted potential interactions between the Krt79^high^ Krt17^high^ follicular cluster and clusters that lay along the rising EpCAM expression axis (monocytes, ISG mono, MCs and resident mLCs). This analysis confirmed that follicular epithelial cells were most likely to signal towards differentiating monocytes (Figure 4G), rather than established mLCs resident within the IL34-rich inter-follicular cells. Assessment of key receptor-ligand interactions identified several potential interactions via *Apoe* from follicular epithelial cells (Figure 4H), consistent with the need for monocytes to adapt to the lipid-rich epidermal environment (Tacke *et al*. 2007). Of potential recruitment pathways, the *Cxcl14*-*Cxcr4* axis was predominantly located to monocytes, in agreement with previous work from the Moser lab showing CXCL14-mediated recruitment of human monocytes to the epidermis before differentiation to mLCs (Schaerli *et al*. 2005). In addition, follicular epithelial cells uniquely provided Jagged-1 and -2 (*Jag1, Jag2*) ligands to initiate Notch signaling in monocytes and, to a lesser extent, MCs (Figures 4H, I). Furthermore, Jag1 expression within EpCAM^+^ keratinocytes was detected at the protein level by flow cytometry (Figure 4J).

**Fig. 4.**
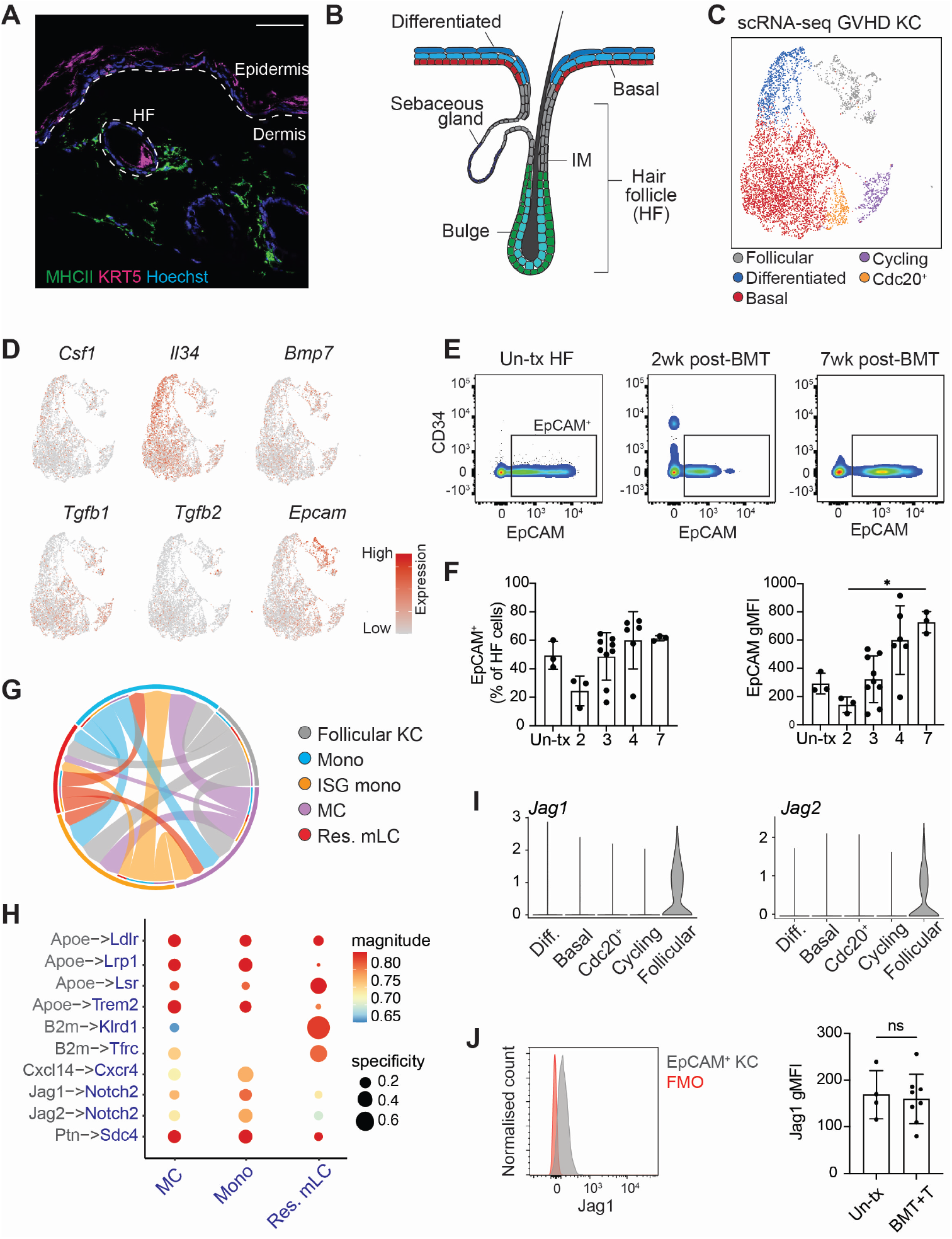
A specialised keratinocyte niche permits mLC differentiation. **A**. Immunofluorescence staining of murine epidermis showing MHCII^+^ (green) cells clustering around KRT5^+^ hair follicle cells (magenta). Hoechst 33342 (blue) was used for nuclear staining. Dotted lines indicate epidermis/dermis border and hair follicle (HF) structure. Scale bar = 50µm. **B**. Schematic representation of hair follicle structure found in the murine epidermis. IM, isthmus. **C**. UMAP visualisation and clustering of keratinocytes from the GVHD analysed by scRNA-seq. KC, keratinocytes **D**. Heatmap overlays showing normalised expression of indicated genes overlaid onto UMAP from (B). **E**. Representative flow plots showing EpCAM expression on hair follicle cells from untransplanted (Un-tx) and indicated time points post-BMT + T cells from epidermis. Gated on live, singlets, CD45^neg^, Sca1^neg^ cells. **F**. Bar graphs showing quantification (let) and gMFI of EpCAM (right) from population indicated in (E) from Un-tx and indicated time points (in weeks) post-BMT+T cells from epidermis. Data are represented as mean±SD (n=3 for Un-tx, n=3 for 2 weeks, 9 for 3 weeks, 7 for 4 weeks, 3 for 7 weeks). Data are pooled from three independent experiments. Significance was calculated using a Kruskal-Wallis test for multiple comparisons, p^*^<0.05. **G**. Chord plot showing the extent of receptor-ligand interactions between follicular KC (grey) and mono (blue), ISG Mono (orange), MC (purple) and res. mLC (red) assessed by LIANA. The width/weight of each arrow indicates the number of potential interactions identified. **H**. Dot plot showing the specificity (NATMI edge specificity) and magnitude (sca LR score) of interactions between follicular KC (grey) and indicated populations (blue). **I**. Violin plots showing normalised expression of indicated genes across scRNA-seq clusters. Diff, differentiated. **J**. Representative histogram (left) of Jag1 expression on EpCAM^+^ CD45^-^ cells from the GVHD epidermis and bar graph (right) showing gMFI of Jag1 from these cells from untransplanted (Un-tx) and GVHD (BMT+T) epidermis. Data are represemted as mean±SD (n=4 for Un-tx, n=8 for BMT+T). Data are pooled from two independent experiments. Significance was calculated using a Mann-Whitney test; ns, not significant.

Our combined data inferred a model in which mLC differentiation was dependent on interactions at distinct epidermal sites; a hair follicle niche to recruit and potentially instruct monocyte differentiation, and an interfollicular niche providing IL34 for survival. Metabolic adaptation of macrophages to utilize fatty acid oxidation pathways is essential for long-term survival as quiescent tissue-resident cells (Wculek *et al*., 2022, 2023). Therefore, we postulated that only differentiated resident mLC would show evidence of metabolic adaptation to the epidermal environment. To test this, we analyzed metabolic pathway usage across cell clusters using COMPASS (Wagner *et al*., 2021), which identifies cellular metabolic states using scRNA-seq data and flux balance analysis (Figure 5A). These data revealed that, despite being present in the epidermis during the same 3-week time-line post-BMT, mLC metabolism was dominated by fatty acid oxidation, whilst the EpCAM^+^ MC expressed higher levels of pathways linked to amino acid metabolism, suggesting active cellular processes. Strikingly, differentiated mLCs also displayed a markedly different metabolic signature to Mrc1^+^ macrophages, despite co-localization in the epidermis. Use of SCENIC (Single Cell Regulatory Network Inference and Clustering) analysis (Aibar *et al*., 2017) to infer transcription factor-target gene groups (regulons) which were more highly active in resident mLCs than other populations, revealed that those regulons enriched within resident mLC were dominated by transcription factors known to be upregulated in response to hypoxia and a lipid rich environment (*Zeb1* or *Rxra* and *Srebf2* respectively) (Figure 5B) (Lengqvist *et al*., 2004; Bommer and MacDougald, 2011; Joseph *et al*., 2015). These data suggested that adaptation to the unique epidermal environment informed mLC development.

**Fig. 5.**
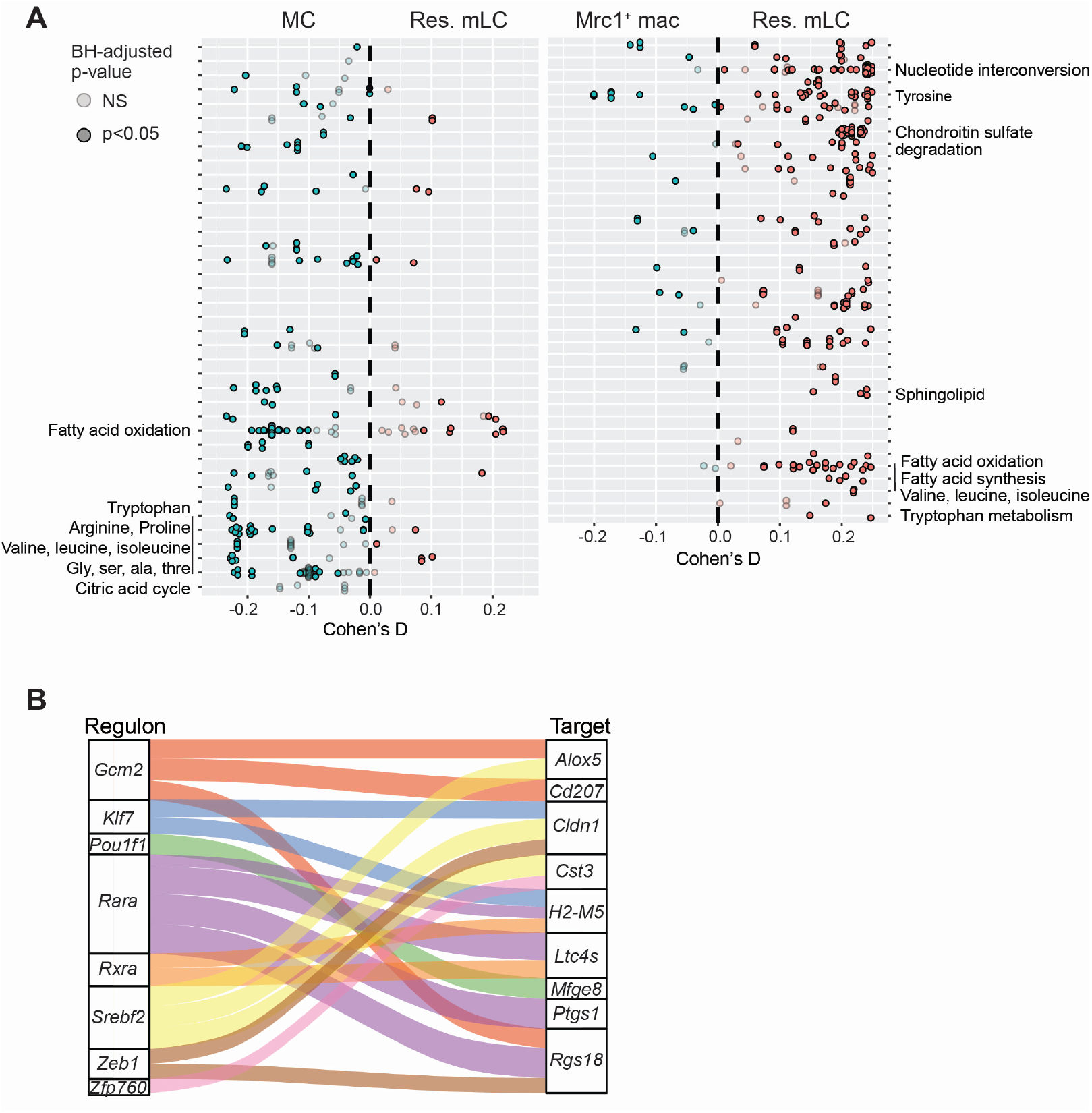
Metabolic adaptation of resident mLC. **A**. Differential activity of metabolic reactions between MC and res. mLC (left) and Mrc1^+^ mac and res. mLC (right) analysed using COMPASS. Reactions (dots) are coloured by the sign of their Cohen’s D statistic, bright dots P<0.05. **B**. Relationship between 8/10 of the most enriched res. mLC regulons identified by SCENIC (Regulon; left) and 9/10 of the res. mLC signature genes shown in Figure S1B (Target; right).

Our findings therefore reveal a precise and spatially restricted hair follicle niche that permits not only recruitment, but potentially commitment of monocytes to resident mLCs in the adult epidermis. This niche is separate from the interfollicular epidermis which provides the IL34 required for maintenance of the differentiated LC network, enabling metabolic adaptation of mLC to the lipid-rich epidermal environment.

### Notch signaling is sufficient to program mLC differentiation

Our data suggested a working model in which recruitment of monocytes to the upper follicular epidermis initiates a molecular cascade resulting in the differentiation of CD207^+^EpCAM^+^ Ahr-expressing mLCs. To define mechanistic pathways leading to a mLC fate in the adult skin we first tested the impact of Notch signaling on monocytes. Provision of Jagged-1 (Jag1), but not the Notch ligand Delta-like 4 (DL4), was sufficient to enhance differentiation of monocytes towards EpCAM^+^ mLC-like cells *in vitro* in the absence of TGFβ and IL34 (Figure 6A). We noted that DL4 appeared to inhibit mLC development in these cultures (Figure 6A). Whilst Notch signaling did not augment mLC frequencies beyond that induced by TGFβ and IL34 (Figure 6A) we observed selection of a mLC-fate at the expense of other default monocyte-derived macrophage- or DC-like cells indicated by expression of CD11c and CD206/CD64. (Figure 6B). Therefore, to determine the effect of Notch signaling combined with other environmental signals in the skin we cultured monocytes with GM-CSF/TGFβ/IL34 with or without Jag1, the Ahr agonist FICZ, or both, and compared the transcriptional changes within sorted CD11b^low^EPCAM^+^ mLC-like cells (Figures 6C and S6A). Figures 6D and E show that cells were clustered by stimulation with considerable overlap between groups. Activation of Ahr signaling did not markedly distinguish clusters along principal component (PC) 1 and over-laid the impact of Jag1, likely due to the dominant activation of the Ahr-responsive gene *Cyp1a1* (Figure 6E, Table S1). However, provision of Jag1 signaling alone led to a marked transcriptional separation suggesting fundamental reprogramming of EpCAM^+^ cells in this group. Comparison to our *in vivo* gene signatures demonstrated that Notch signaling was sufficient to program mLCs that expressed both LC-defining transcription factors (*Id2, Ahr*) and genes upregulated within the skin environment (*Epcam, Cd207, Cldn1, Mfge8*) (Figure 6F). Moreover, direct comparison of our scRNA-seq gene signatures defining *in vivo* epidermal cell populations within our bulk RNA-seq dataset demonstrated that Jag1 signaling prescribed resident mLC identity (Figure 6G). The additional activation of Ahr signaling pushed cells towards a cycling mLC phenotype (Figure 6G).

**Fig. 6.**
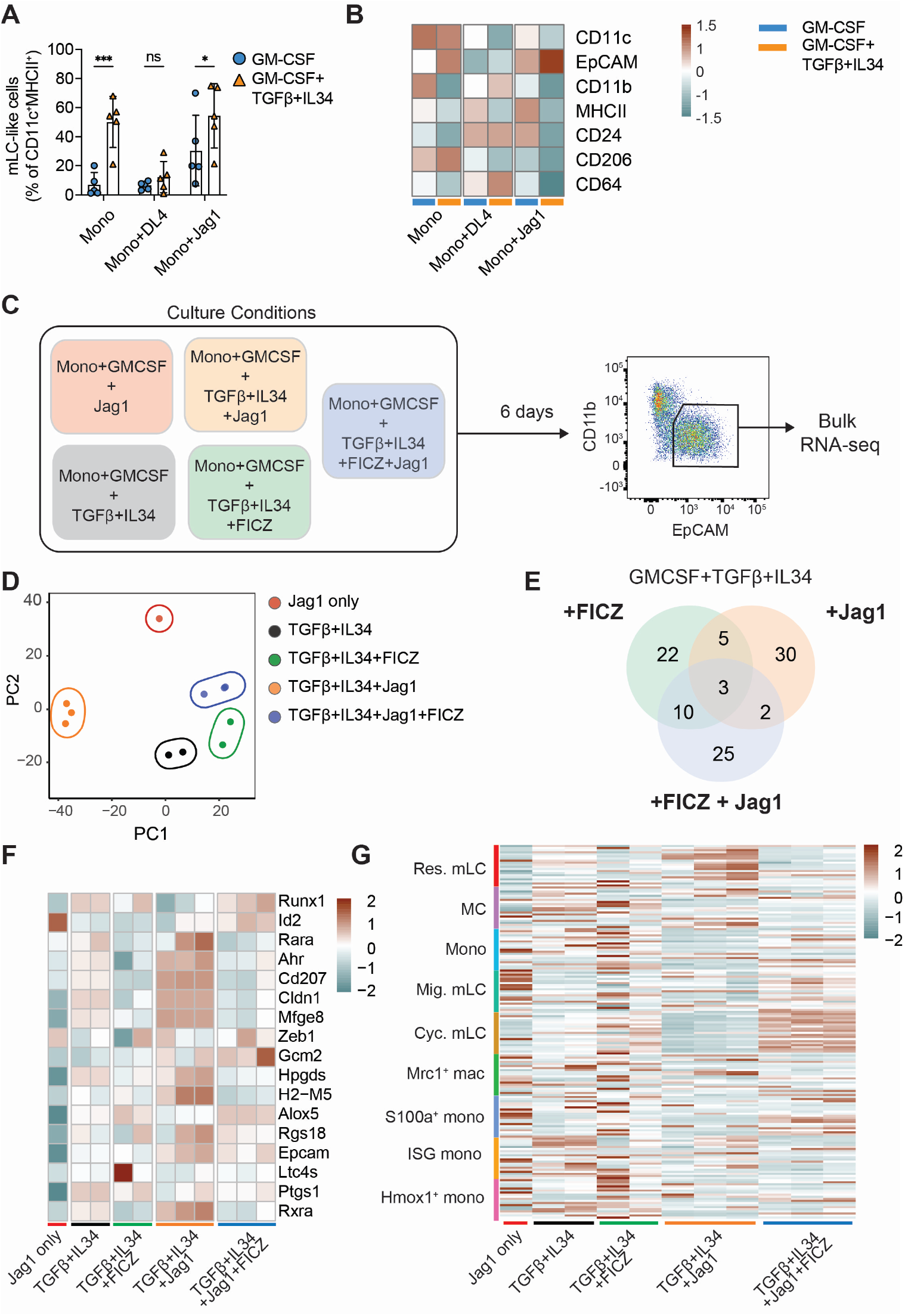
Notch signaling programs mLC differentiation. **A**. Bar graph showing the proportion of mLC-like cells generated from monocytes cultured with GM-CSF alone or GM-CSF, TGFβ and IL34 in the presence or absence of indicated Notch ligands (see figure S6A for gating strategy). Data is shown as mean±SD (n=5). Significance was calculated by 2 way ANOVA with uncorrected Fisher’s LSD for multiple comparisons, ^*^ p<0.05; ^***^p<0.001. **B**. Heatmap showing average gMFI of indicated markers from bone-marrow derived monocytes cultured and analysed by flow cytometry as indicated in (A) (n=5). **C**. Experimental set up for bulk RNA-seq of mLC-like cells generated under indicated conditions. **D**. PCA plot of bulk RNA-seq samples coloured by culture condition. **E**. Venn diagram showing numbers of common and unique DEGs between indicated conditions. **F**. Heatmap showing scaled expression of LC signature genes across samples. **G**. Heatmap showing expression of gene signatures from epidermal myeloid cell clusters (defined as top 20 DEGs) (y-axis) across bulk RNA-seq samples (x-axis).

Thus, Notch signaling is sufficient to restrict the differentiation potential of monocyte-derived cells, directing them away from a default macrophage program towards a mLC fate.

### Post-natal maturation of eLCs in human skin induces expression of DC-like immune gene programs that mirror mLC development

Our data demonstrate that MDP-derived monocytes adopt a distinct pathway of differentiation in murine adult skin, characterised by loss of Zeb2, to become mLCs. We therefore questioned whether such a transition away from a classical macrophage identity occurred in humans. Post-natal maturation of intestinal macrophages has been linked to the acquisition of immune functions (Viola *et al*. 2023). We posited that a similar maturation process occurred in human skin after birth, and was linked to LC specification. To address this, we analyzed a unique collection of samples taken from newborn babies (<28 days), infants (1 month - 1 year) and children (2-15 years) (Figure 7A) by bulk RNA-seq. As previously demonstrated in mice and humans, we observed a marked expansion in eLC numbers in the transition from newborns to infants (Figure 7B). Visualisation of the RNA-seq data as a co-expression network revealed a central gene program that is high in newborns and encodes basic macrophage functions including protein transport, RNA processing, and cadherin binding (Cluster 1, 2, and 4, Figure 7C and Table S2). Specifically, there was a significant increase in transcriptional activity associated with induced immune activation from newborns to infants and children (Cluster 5, Figure 7C,D) that was also enriched for biological processes such as antigen processing and presentation (Figures 7E and Table S2), suggesting that more DC-like functions were activated once the skin environment and LC network has fully matured. Furthermore, expansion of cluster 5 was associated with a significant increase in transcriptional activity (Figure 7C).To determine how post-natal maturation of eLCs in human skin compared to mLC differentiation in adult murine skin we analyzed expression of the key factors associated with mLC development. Proliferation of eLCs in infant skin resulted in a marked loss of ZEB2 expression with concomitant upregulation of EPCAM, AHR and AHRR (Figure 7F).

**Fig. 7.**
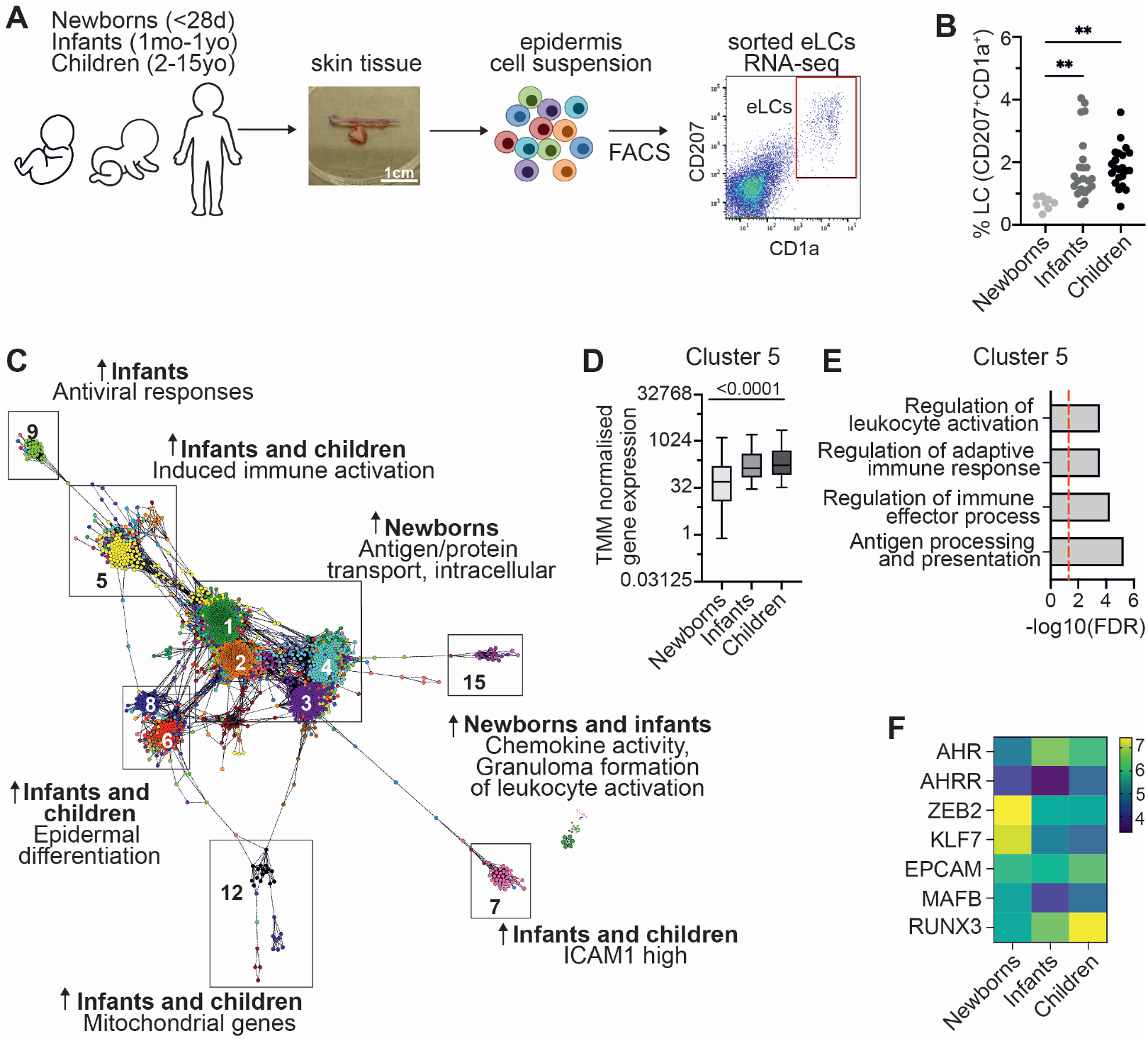
Post-natal maturation of eLCs in human skin induces expression of DC-like immune gene programs that mirror mLC development. **A**. Human LCs isolation workflow. Skin samples were collected from healthy donors aged 0-15 yo and epidermal cell suspensions were obtained. CD207^+^CD1a^+^ cells were FACS purified directly into Trizol. d, days; mo, months; yo, years old. **B**. Percentage of CD207^+^CD1a^+^ cells across newborns, infants and children. Significance was calculated by one way ANOVA, ^**^p<0.01. **C**. Average TMM normalised gene expression levels in cluster 5 across newborns, infants and children. Significance was calculating using one way ANOVA. **D**. Transcript to transcript clustering Graphia, 2447 genes, r=0.75, MCL=1.7 identified 21 clusters with n>10 genes, encoding distinct transcriptional programs in human LCs. Arrows indicate enrichment. **E**. Gene ontology ranked with FDR corrected p-values given for cluster 5. **F**. Heatmap showing normalised expression of indicated genes.

In summary, we demonstrate that monocytes recruited to the epidermis after immune pathology adopt a unique pathway of differentiation to mLCs that mirrors post-natal program in human skin.

## Discussion

Whether LCs are macrophages or DCs has long been debated (Lutz *et al*. 2017) and both denominations are still routinely applied in the literature, despite fate mapping studies demonstrating the embryonic macrophage origin for these cells (Ginhoux *et al*. 2006; Hoeffel *et al*. 2012; Schulz *et al*. 2012; Mass *et al*. 2016). Here we have begun to resolve this discussion demonstrating a selective bias in the cells that replenish the LC network after destruction of eLCs, whereby MDP-derived monocytes are more likely to become mLCs. Within the skin, localisation and signaling within a unique hair follicle niche drives loss of Zeb2, the regulator of tissue-resident macrophage identity at other barrier sites, and Notch-dependent expression of the LC-defining transcription factors Id2 and Ahr, to generate long-lived mLCs (Chopin *et al*. 2013; Mass *et al*. 2016). This adaptation to adult skin mirrors post-natal maturation of eLCs which is characterised by the appearance of a specific gene program associated with DC-like immune functions. Thus, specification of LCs within the skin environment drives evolution of a unique population of tissue-resident macrophages, which are the only resident macrophages that can migrate to draining lymph nodes and prime T cell immunity.

Inflammation due to UV irradiation or GVHD leads to influx of wave(s) of monocytes, the majority of which will not succeed in becoming long-lived LCs (Ginhoux *et al*., 2006; Seré *et al*., 2012). We previously used mathematical modelling to define the cellular processes by which monocytes refill the empty epidermis, demonstrating a significant inefficiency whereby only 4% of epidermal monocytes were predicted to become long-lived mLCs (Ferrer *et al*., 2019). Our data now explain this inefficiency by demonstrating: first, that only a subset of Ly6C^+^ monocytes can become mLCs; and second, that LC precursors compete for Notch signals within a spatially restricted hair follicle niche. There is a growing awareness of the heterogeneity within Ly6C^+^ monocytes, that represent a pool of cells derived from either GMPs or MDPs (Yáñez *et al*., 2017; Kwok *et al*., 2020; Trzebanski *et al*., 2023). Here we show that, despite the relative scarcity of MDP compared to GMP in BM, MDP-Mos selectively accumulate in the epidermis to become mLCs. MDPs were also more efficient than GMPs in giving rise to CD11b^+^EpCAM^+^ mLCs *in vitro*, suggesting intrinsic bias towards responsiveness to epidermal growth factors. Further experiments are needed to test whether the relative abundance of MDP-Mos in the epidermis is due to their selective recruitment and/or preferential expansion and survival of these cells within the epidermal niche.

Evidence for the macrophage origin of embryonic LCs comes from reporter studies in which labelled yolk sac (Schulz *et al*., 2012; Mass *et al*., 2016) and fetal liver (Hoeffel *et al*., 2012) macrophage/monocytes gave rise to the nascent LC network in the developing embryo. Moreover, use of the macrophage-restricting gene MafB to lineage trace myeloid cells labelled all LCs in the healthy adult skin (Wu *et al*., 2016), and generation of LCs from CD34^+^ stem cells requires suppression of the transcription factor Klf4, which is associated with differentiation of moDCs (Jurkin *et al*., 2017). But, LCs express high levels of CD24 and Zbtb46, considered markers of conventional DCs (Wu *et al*., 2016), and activation and migration initiates the expression of a convergent transcriptional program shared with DC populations leaving the skin (Clayton *et al*., 2017; Polak and Singh, 2021; Reynolds *et al*., 2021). Migrating LCs upregulate an IRF4-dependent gene program which is also evident in moDC (Sirvent *et al*., 2020; Davies *et al*., 2022). We believe our data begin to reconcile this dichotomy. Zeb2 is a critical regulator of cell fate that specifies tissue-resident macrophage identity across barrier sites (Scott *et al*., 2018; Anderson *et al*., 2021) and we show that only those cells that have lost Zeb2 expression upregulate EpCAM to become mLCs. Key regulatory elements control function in embryonic and HSC-derived macrophages (Huang *et al*., 2021) and molecular cross-talk with Id2 has been shown to determine cDC2 specification (Liu *et al*., 2022). We speculate that a similar process may permit expression of Id2 in differentiating mLCs and initiating the shift from more macrophage-like to DC-like cells.

To determine the extrinsic signals regulating mLC development we characterized the epidermal niche and identified a spatially restricted area of the follicular epidermis at which homotypic binding of EpCAM^+^ MCs provides access to local Notch signals. The epidermis is composed of layers of stratified epithelial cells which are interspersed in non-glabrous skin with follicular structures that support the cycles of hair growth. Within follicular epithelial cells tightly demarcated areas of CCL2 expression at the upper follicular isthmus is associated with recruitment of monocytes to the epidermis (Nagao *et al*., 2012), which is spatially distinct from sites of integrin αvβ6 expression required for activation of latent TGFβ and retention of mature LCs in the interfollicular epidermis (Mohammed *et al*., 2016). MHCII^+^CD11b^+^ cells accumulated around hair follicles upon induction of inflammation in LC-depleted bone marrow (BM) chimeras, and LC repopulation was impaired in mice lacking hair follicles (Nagao *et al*., 2012). But whether the hair follicle site provided a differentiation niche, or merely served as a point of entry for monocytes into the epidermis was not known. Our data suggest that the follicular niche is not only a site of recruitment but also provides instructive signals via Notch signaling for differentiation to mLCs before they relocate within the keratinocytes of the differentiated epidermis. Our findings reflect the importance of Notch for differentiation of monocyte-derived Kupffer cells in the liver (Bonndardel *et al*) hinting at shared exploitation of Notch pathways across tissue macrophage niches, although we identified Jagged as the ligand rather than DLL4 in the liver (Bonnardel *et al*., 2019). Unlike the liver, however, we suggest that the physical restraints in the epidermis and separation from the circulation impose a two niche model whereby hair follicle keratinocytes instruct LC molecular identity, but the differentiated intra-follicular keratinocytes provide the scaffold and trophic factors that support survival and permit adaptation to the lipid-rich epidermal environment.

Notch signaling has previously been linked to human moLC development *in vitro* (Strobl, Krump and Borek, 2019; Bellmann *et al*., 2021), while a recent study that employed scRNA-seq of human LCs revealed the presence of 2 eLC populations in the skin, linking Notch signaling to expansion of EpCAM^neg^ cells (Liu *et al*., 2021). By contrast, our data suggest that Notch signaling restricts murine monocyte differentiation into mLCs at the expense of other fates. It is possible that Notch signaling promotes a more DC-like program in mLCs; Notch signaling in monocytes has been shown to suppress a macrophage fate in favor of a DC fate and is required for differentiation to monocyte-derived CD207^+^CD1a^+^ cells characteristic of LC histiocytosis (Kvedaraite *et al*., 2022), while activation of Notch in human CD1c^+^ DCs is sufficient to promote differentiation to LC-like cells that contain Birbeck granules (Milne *et al*., 2015).

Ahr is an evolutionarily conserved cytosolic sensor that functions as a ligand-dependent transcription factor to control cell fate decisions in gut immune cells (Stockinger, 2009), and direct monocytes towards a DC-like rather than a macrophage endpoint (Goudot *et al*. 2017). The role of Ahr signaling in LCs remains unclear; eLC begin to express Ahr upon differentiation *in utero* (Mass *et al*. 2016), but our data suggest that expression increases post-birth. The epidermis of Ahr-deficient mice is replete with LCs (Esser, Rannug and Stockinger, 2009), albeit a less activated population, probably due to the absence of dendritic epidermal T cells (DETC) and reduced GM-CSF production in the skin of these mice (Jux, Kadow and Esser, 2009). Moreover, mice fed with chow deficient in Ahr dietary ligands had normal numbers of LCs (Cros *et al*. 2023), but these cells did not migrate to draining LNs. In contrast, in one study, LC-specific deletion of Ahr led to a reduction in epidermal LCs (Hong *et al*. 2020). We tested the role of Ahr signaling for the differentiation of mLCs and found that blockade of Ahr prevented monocyte differentiation *in vitr*o, supporting a previous study using CD34^+^ precursors (Platzer *et al*. 2009). However, use of competitive chimeras demonstrated that Ahr-deficient monocytes could become mLCs *in vivo*. We observed that canonical (Cyp1b1) Ahr signaling was active in mLCs *in vitro* but not *in vivo* suggesting activation of alternative pathways within the epidermal environment.

Murine and human eLCs undergo a burst of proliferation after birth (Chorro *et al*., 2009; Meindl *et al*., 2009), but it was not known whether expansion of eLC was associated with maturation of the network, as has been demonstrated for the LCs of the oral mucosa (Jaber *et al*., 2023). To address this question, we assembled a unique RNA-seq dataset from eLCs sorted from newborn children, up to 1 year-old infants and older children. These data revealed the documented increase in LC density after the first 28 days of life (Meindl *et al*., 2009) and demonstrated that this increase was associated with a marked change in gene expression whereby the macrophage-associated genes MAFB and ZEB2 were down-regulated, while we observed an increase in AHR and EP-CAM. Notably, this transition was accompanied by the expression of gene modules associated with enhanced immune and DC-like functions. These findings support our proposed concept of gene regulatory networks defining LC function whereby interaction between Ahr and Irf4 activates expression of immunogenic function and migration to LNs (Polak and Singh, 2021). The signals that trigger eLC proliferation in the skin are not known. Whilst skin commensal bacteria per se are not required for LC development and survival (Capucha *et al*., 2018), it is possible that the increase in microbiota diversity during the first year of life could play an important role in conditioning the LC niche. Further experiments are required to test this hypothesis.

In conclusion, convergent evolution of monocytes within adult skin imposes expression of these gene programs to mirror post-natal conditioning and maintain this unique population of DC-like cells in the epidermis.

## Materials and methods

### Mice

C57BL/6 (B6) were purchased from Charles River UK. Langerin.DTR^GFP^ were originally provided by Adrian Kissenpfennig and Bernard Malissen (Kissenpfennig *et al*., 2005), T cell receptor (TCR)-transgenic anti-HY Matahari were provided by Jian Chai (Imperial College London, London, UK)(Valujskikh *et al*., 2002) and CD45.1 mice were bred in-house at University College London (UCL) Biological Services Unit. ID2^BFP^ reporter mice were a kind gift from Andrew McKenzie (University of Cambridge), whereby the pBAD-mTagBFP2 plasmid was a gift from Vladislav Verkhusha (Addgene plasmid no. 34632 ; http://n2t.net/addgene:34632 ; RRID:Addgene_34632) (Subach *et al*., 2011; Walker *et al*., 2019). All procedures were conducted in accordance with the UK Home Office Animals (Scientific Procedures) Act of 1986 and were approved by the Ethics and Welfare Committee of the Comparative Biology Unit (UCL, London, UK).

### Human samples

Human skin samples were collected with written informed consent from donors with approval by the South East Coast - Brighton & Sussex Research Ethics Committee in adherence to Helsinki Guidelines (ethical approvals: REC approval: 16/LO/0999).

### Bone marrow transplants

Recipient male CD45.2 B6 mice were lethally irradiated (10.4Gy of total body irradiation, split into two fractions separated by 48 hours) and reconstituted 4 hours after the second dose with 5 x 10^6^ female CD45.1 B6 bone marrow (BM) cells, 2 x 10^6^ CD4 T cells, with 1 x 10^6^ CD8 Matahari T cells administered by intravenous injection through the tail vein. CD4 and CD8 donor T cells were isolated from spleen and lymph node single cell suspensions by magnetic activation cell sorting (MACS; Miltenyi) using CD4 (L3T4) and CD8a (Ly-2) microbeads (Miltenyi) according to manufacturer’s instructions. In some experiments, BM from ID2^BFP^ B6 mice was used to track donor LC. In some experiments, Langerin.DTR^GFP^ male mice were used as recipients to track host LCs.

### Mixed chimera experiments

BM from AHR^-/-^ mice was a gift from Brigitta Stockinger (Francis Crick Institute) (Schmidt *et al*., 1996). Lethally irradiated male Langerin.DTR^GFP^ B6 mice received a 50:50 mix of BM from AHR^-/-^ and ID2^BFP^ female mice, with CD4 T cells and CD8 Matahari T cells. Three weeks following transplant, epidermis and spleens were processed and analysed for chimerism by flow cytometry.

### Tissue processing. Murine skin

Epidermal single cell suspensions were generated as described (Ferrer *et al*., 2019; West *et al*., 2022). Dorsal and ventral sides of the ear pinna were split using forceps. These were floated on Dispase II (2.5mg/ml; Roche), in HBSS and 2% FCS for 1 hour at 37°C or overnight at 4°C, followed by mechanical dissociation of the epidermal layer by mincing with scalpels. Cells were passed sequentially through 70- and 40µm cell strainers in 1mM EDTA, 1% FCS, PBS solution. **Human skin:** Fat and lower dermis was cut away and discarded before dispase (2 U/ml, Gibco) digestion for 20h at 4°C. Epidermal sheets were digested in Liberase™ (13 U/ml, Roche), for 1.5h at 37°C. **Bone marrow:** Bone marrow single cell suspensions were prepared from femurs and tibias of donors using a mortar and pestle. Red blood cells were lysed in 1ml of ammonium chloride (ACK lysis buffer; Sigma) for 1 min at room temperature. Cells were washed and resuspended in complete RPMI (RPMI supplemented with 10% FCS, 1% L-glutamine, 1% Penicillin-streptomycin) until used.

### *In vitro* cultures. GMP and MDP cultures

GMP, MDP and Ly6C^hi^ monocytes from whole bone marrow were FACS isolated and up to 2 x 10^3^ cells were seeded in 96-well flat-bottom tissue-culture treated plates. Cells were cultured in complete RPMI and supplemented with with recombinant GM-CSF (20ng/ml; Peprotech), TGFβ (5ng/ml; R&D systems) and IL34 (8µg/ml; R&D systems). Cells were cultured at 37°C, 5% CO_2_. The medium was partially replaced on day 2 of culture and completely replaced on day 3, and cells were harvested on day 6. **Monocyte cultures:** Monocytes were isolated from whole bone marrow by MACS using Monocyte Isolation Kit (BM; Miltenyi, 130-100-629) as per manufacturer’s instructions. Monocytes were resuspended in complete RPMI and plated at 5 x 10^5^ cells per well in tissue-culture treated 24-well plates. Cells were cultured as described above. **Monocyte-OP9 co-cultures:** OP9, OP9-Jag1 and OP9-DL4 cell lines were gifted by Victor Tybulewicz (Francis Crick Institute) and were cultured in RPMI supplemented with 10% FCS, 1% L-glutamine, 1% Penicillin-streptomycin, MEM NEAA, sodium pyruvate, HEPES buffer and ß-mercaptoethanol. OP9 cells were seeded onto tissue-culture treated 24-well plates at 2 x 10^4^ cells per well and incubated overnight at 37°C. The next day, following monocyte isolation, cells were counted, and 1 x 10^5^ monocytes were seeded on to OP9 cells. Cells were cultured as described above.

### Flow cytometry and cell sorting. Murine

Cells were distributed into 96-well V-bottom plates or FACS tubes and incubated in 2.4G2 hybridoma supernatant for 10 min at 4°C to block Fc receptors. Cells were washed with FACS buffer (2% FCS, 2mM EDTA, PBS) before adding antibody cocktails that were prepared in a total volume of 50µl per test in Brilliant Stain Buffer (BD Biosciences) and FACS buffer. Cells were incubated with antibodies for 30 mins on ice then washed with FACS buffer. Viability was assessed either by staining cells with Fixable Viability Dye eFluor680 (eBiosciences) or Propidium iodide (PI), for fixed or unfixed cells respectively. For intracellular staining, cells were fixed and permeabilised using the Foxp3/Transcription Factor Staining Buffer Set (Invitrogen) for 30 mins on ice. Cells were subsequently washed in permeabilisation buffer before adding antibody cocktails that were prepared in a total volume of 50µl per test in permeabilisation buffer. Cells were incubated with antibodies for 30 mins on ice then washed with permeabilisation buffer. Antibodies used are listed in table S3. **Human:** Antibodies used for cell staining were pre-titrated and used at optimal concentrations. For FACS purification LCs were stained for CD207 (anti-CD207 PeVio700), CD1a (anti-CD1a VioBlue) and HLA-DR (anti-HLA-DR Viogreen, Miltenyi Biotech, UK).

When required, cells were acquired on a BD Fortessa analyzer equipped with BD FACSDiva software; or sorted into either complete RPMI or RLT lysis buffer (Qiagen) or Trizol using a BD Aria III.

### Single cell RNA-sequencing

Single cell suspensions from murine epidermis were stained for FACS as described above. Donor and host CD11b^+^MHCII^+^ cells as well as CD45^neg^ cells were sorted into RPMI medium supplemented 2% FCS and counted manually. Cell concentrations were adjusted to 500-1200 cells/µl and loaded at 7,000-15,000 cells per chip position using the 10X Chromium Single cell 5’ Library, Gel Bead and Multiplex Kit and Chip Kit (10X Genomics, V3 barcoding chemistry) according to manufacturer’s instructions. Data are from multiple mice sorted from 2 independent BMT experiments. All subsequent steps were performed following standard manufacturer’s instructions. Purified libraries were analysed by an Illumina Hiseq X Ten sequencer with 150-bp paired-end reads. ***scRNA-seq data processing and analyses***. Generated scRNA-seq data were preprocessed using the kallisto and bustools workflow (Melsted *et al*., 2021). Downstream analysis was performed using the Seurat package in R (Stuart *et al*., 2019). Cells with <500 detected genes and >20% mitochondrial gene expression were removed from the dataset. DoubletFinder was used to identify and remove any likely doublets. These were typically less than 1% of each batch. PCA was performed on the 2000 most variable genes and clusters were identified using the Leiden algorithm. Clusters were annotated based on the expression of key cell-type defining genes. Differentially expressed genes (DEGs) were identified using the FindMarkers function with significance cut offs of log2-fold change > 2 and adjusted p-values < 0.05. ***Enrichment score analysis***. To calculate enrichment scores for specific gene signatures the Seurat function AddModuleScore was used. The human migLC gene signature included 101 genes (Sirvent *et al*., 2020). The MDP-Mo and GMP-Mo signatures included 140 and 108 genes respectively (Trzebanski *et al*., 2023). ***Trajectory analyses***. Pseudotime trajectory inference of differentiation and RNA velocity analysis based upon spliced and unspliced transcript ratios were performed respectively using the Slingshot (Street *et al*., 2018) and velociraptor packages for R (La Manno *et al*., 2018). Expression of genes changing along the trajectories were identified using general additive models fitted by tradeseq. ***Receptor-ligand interaction analysis***. Potential receptor-ligand (R-L) interactions between the follicular keratinocyte subset and monocyte-derived cell clusters were investigated using the LIANA package (Dimitrov *et al*., 2022). LIANA is an umbrella framework which creates a consensus R-L score from the methods and pathway tools of several other software packages. It encompasses CellPhoneDB (v2), CellChat, NATMI, iTALK and CytoTalk (Dimitrov *et al*., 2022). ***SCENIC analysis***. SCENIC was used to identify regulons, sets of transcription factors and their cofactors co-expressed with their downstream targets in single cell data (Aibar *et al*., 2017). This analysis was applied to the 10X scRNA-seq dataset and each of the regulon AUC per cell scores was used to identify regulons with the greatest mean difference between Res-mLC and all other clusters. ***COMPASS analysis***. *In silico* flux balance analysis was conducted via COMPASS (Wagner *et al*., 2021). Normalised scRNA-seq counts per million (CPM) gene expression profiles were exported and COMPASS analysis was conducted using standard settings on a high-performance computing cluster (Butcher, King and Zalewski, 2017). Metabolic reactions were mapped to RECON2 reaction metadata (Thiele *et al*., 2013), and reaction activity scores were calculated from reaction penalties. Reactions which do not have an enzyme commission number or for which there is no biochemical support (RECON2 confidence score = 1-3) were excluded from the analysis. Differential reaction activities were analysed via Wilcoxon rank-sum testing and resulting p-values were adjusted via the Benjamini-Hochberg (BH) method. Reactions with an adjusted p-value of less than 0.1 were considered differentially active. Effect sizes were assessed with Cohen’s d statistic.

### Bulk RNA-seq and analyses

Up to 1 x 10^5^ cells were FACS sorted directly into RLT lysis buffer (Qiagen) supplemented with 14mM β-mercaptoethanol. Cells were vortexed immediately after being sorted to ensure cell lysis. RNA was extracted using the RNeasy Micro kit (Qiagen) as per manufacturer’s instructions, with an additional DNA clean-up step using RNase-free DNase I (Qiagen). RNA quantification and quality check was carried out by Novogene (UK), as well as subsequent library preparation and sequencing. Sequencing was performed on a NovaSeq 600 System (Illumina) to yield an average of 30 million reads per sample.

RNA-seq transcript abundance was quantified using the salmon read mapper and an Ensembl GRCm39 transcript model. The data were imported to the R statistical environment and summarised at the gene level (that is, transcript counts summed) using tximport. Statistical transformations for visualisation (vst and log10) and analyses of differential expression were performed using the DESeq2 package (Love, Huber and Anders, 2014). Multiple testing adjustments of differential expression utilised the Benjamini–Hochberg false discovery rate (fdr).

### RT-PCR

RNA was extracted from samples as described above. RNA quantification was carried out using a Nanodrop and cDNA was synthesised using the High-Capacity cDNA Reverse Transcription Kit (Applied Biosystems) as per manufacturer’s instructions. RT-PCR was run on a QuantStudio 5 Real-Time PCR system (Thermo Fisher Scientific) using the Maxima SYBR Green/ROX qPCR Master Mix (2X) (Thermo Fisher Scientific) according to manufacturer’s instructions. Primers used in this study: *Ahr* forward, AGC CGG TGC AGA AAA CAG TAA; *Ahr* reverse, AGG CGG TCT AAC TCT GTG TTC; *Cyp1b1* forward, ACG ACG ATG CGG AGT TCC TA; *Cyp1b1* reverse, CGG GTT GGG AAA TAG CTG C; *GAPDH* forward, CGGGTTCC-TATAAATACGGACTGC; *GAPDH* reverse, GTTCACACC-GACCTTCACCA.

### Immunofluorescence imaging

Skin biopsies were embedded in OCT compound (Leica). 10 mm sections were cut using a cryostat (Leica) and stored at -20°C. Tissue was blocked for 2 hours at room temperature with 5% BSA (Sigma Aldrich), 5% Donkey Serum (Merck) in 0.01% PBS-Tween-20 (PBST). Sections were incubated with primary antibodies for KRT5 (Covance; PRB-160P) and MHCII (Abcam; ab15630) overnight at 4°C. Antibodies were detected using donkey Cy2- or Cy5-conjugated secondary Fab fragment antibodies (Jackson Laboratories) and nuclei stained using Hoechst 33342 (1:1000, Sigma Aldrich), and mounted using Prolong Gold anti-fade mounting media. Images were acquired on a Leica SP8 confocal microscope and subsequently analyzed using NIH ImageJ software.

### Statistical analyses

All data, apart from RNA-seq data, were analyzed using GraphPad Prism Version 6.00 for Mac OsX (GraphPad Software, USA). All line graphs and bar charts are shown as mean±SD. Protein expression data for flow cytometry is shown as geometric mean fluorescent intensity (as specified in figure legends) with the range. Significant differences were determined using one-way analysis of variance (ANOVA) to measure a single variable in three groups or two-way ANOVA for experiments with more than one variable, with post-tests specified in individual figure legends. For comparisons between 2 paired groups a paired t test was used according to a normality test. Significance was defined as ^*^p < 0.05, ^**^p < 0.01, ^***^p < 0.001, ^****^p <0.0001. Statistical details of the data can be found in each figure legend. Analysis of bulk and scRNA-seq data was performed in the R and Python environments using tests described in the method details.

## Data availability

Sequencing data is available for review on request at NCBI GEO GSE247879 and GSE251705.

## Acknowledgments

We thank the UCL Biological Services for their support with animal work. We are grateful to Carlos Minutti and Caetano Reis e Sousa support with reagents and experimental advice. This study was funded by Biotechnology and Biological Sciences Research Council grant BB/T005246/1 (A.A., J.D. and C.L.B.). Acquisition and analysis of human perinatal samples, MEP, SS were funded by the Wellcome Trust Sir Henry Dale Fellowship 109377/Z/15/Z.

## Author contributions

Conceptualization, C.L.B., M.E.P.; Methodology, A.A., J.D., S.S., S.H., S.T., J.S., M.L.L., I.B.C., S.M.H, S.L, E.E. and S.J; Investigation, A.A., J.D., S.S.; Formal analysis, S.S., S.H. and M.E.P, A. V; Resources: S.T, C.M., A. V., N.J.H., M. A-J., S.J; Writing – Original Draft, C.L.B and M.E.P; Writing - Review and Editing, A.A., S.H., S.M.H, E.E., S. J., M. A-J, M.E.P and C.L.B; Funding Acquisition and supervision, C.L.B and M.E.P.

## Conflicts of interest

MEP is currently employed at Johnson and Johnson Innovative Medicine. Johnson and Johnson Innovative Medicine or any of employees/stakeholders have not been involved in any part or aspect of the project or manuscript.

Data and materials are available upon request.

