## Supplemental Information for "Convergent evolution of monocyte differentiation in adult skin instructs Langerhans cell identity"

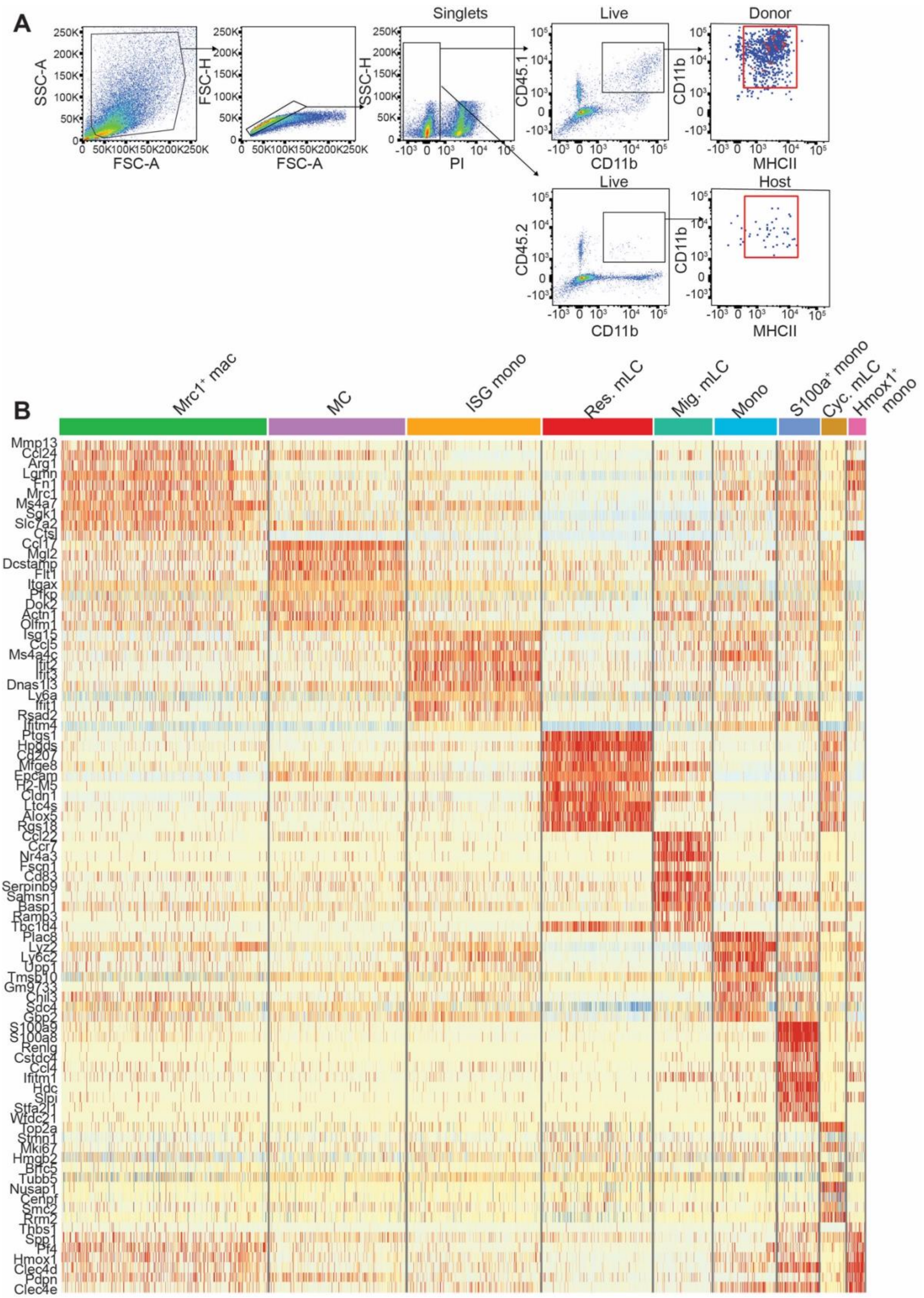


**Figure S1. scRNA-seq of monocyte-derived cells in the inflamed epidermis, related to Figure 1. A**. Gating strategy for FACS isolation of donor and host CD11b⁺MHCII⁺ cells from murine epidermis for 10X scRNA-seq. **B.** Heatmap showing differentially expressed genes across clusters from dataset in figure 1B.

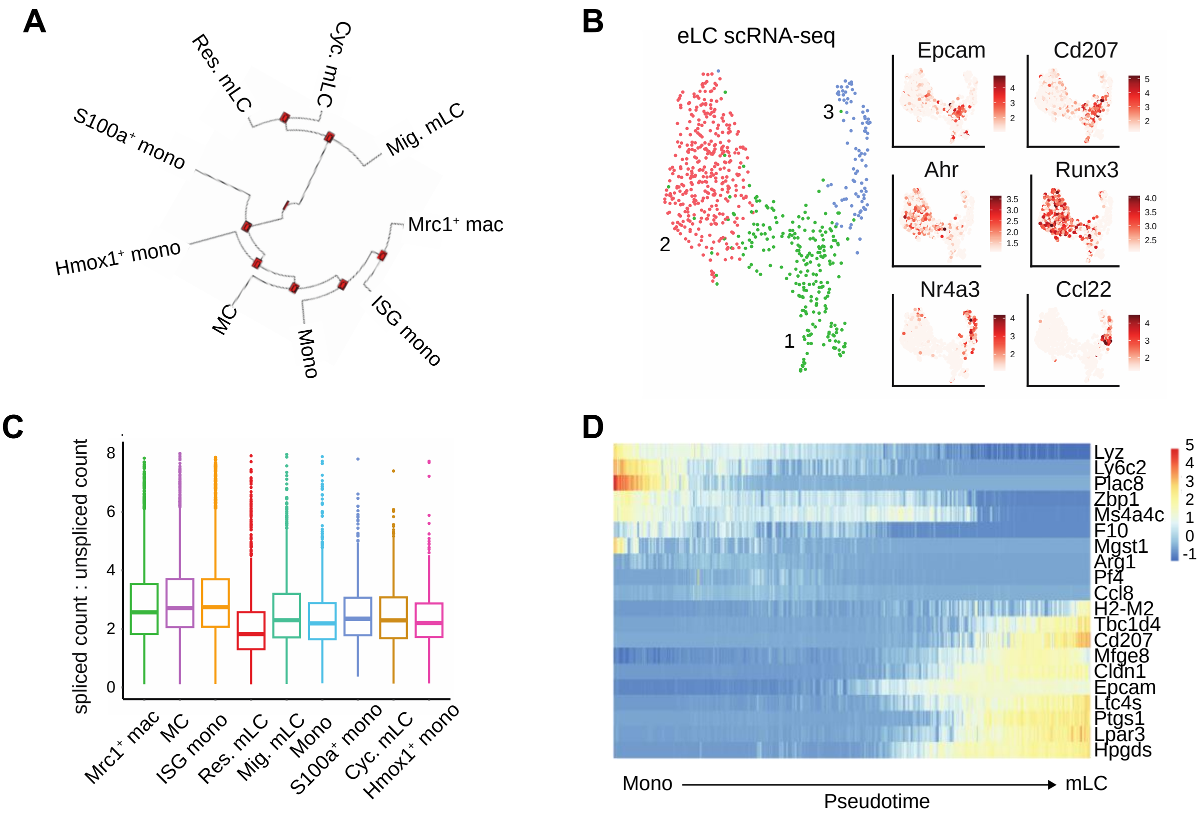


**Figure S2. scRNA-seq of myeloid cells in the inflamed epidermis, related to Figure 1. A.** Parametric bootstrap showing single cell significance of hierarchical clustering identified in Figure 1B. **B.** UMAP and clustering of host LC (left) and heatmap overlays showing expression of indicated genes (right). **C.** Boxplot showing ratio of spliced to unspliced counts across indicated clusters. **D.** Heatmap showing the change in expression of indicated genes across pseudotime from monocyte to mLC differentiation.

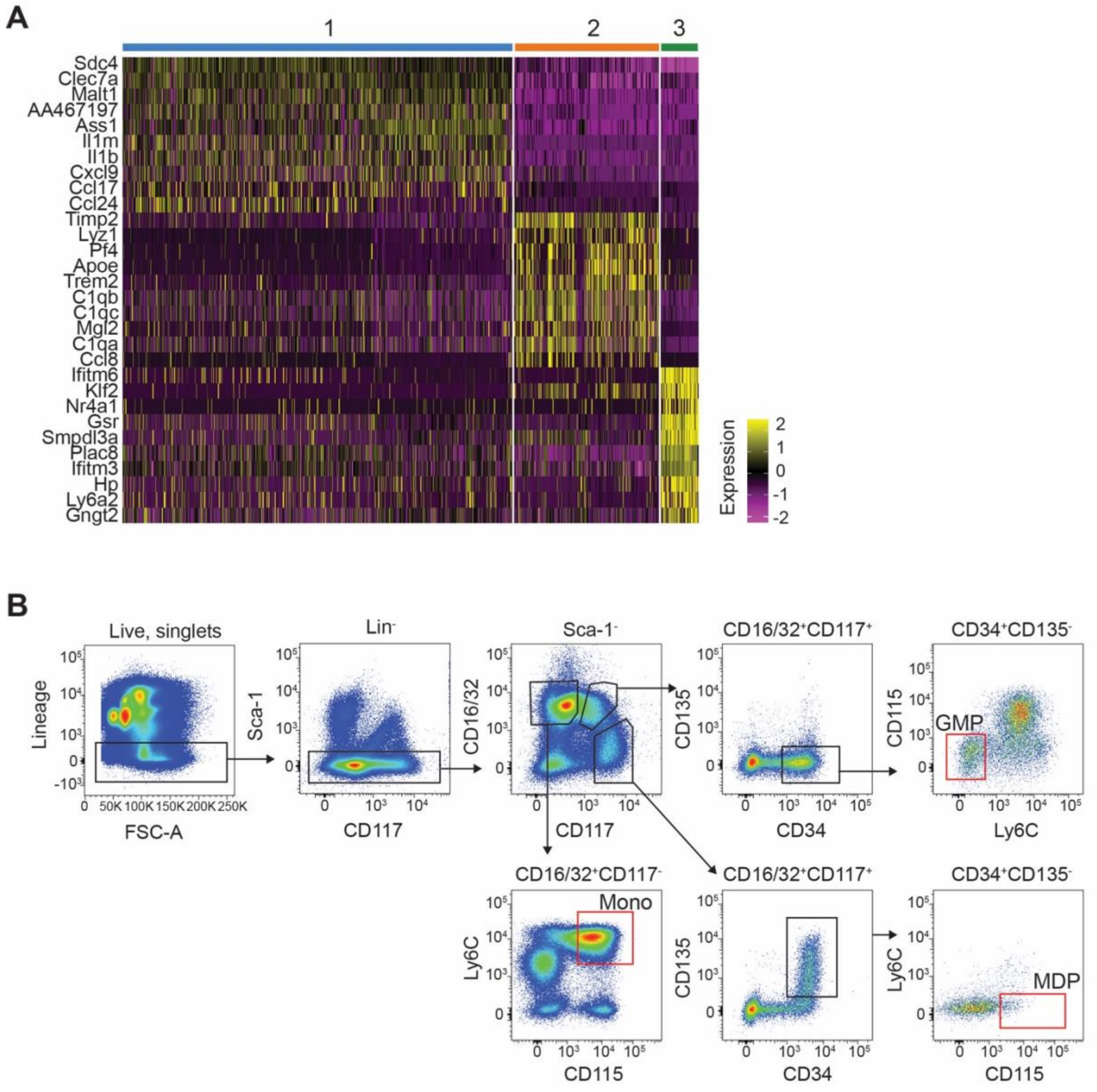


**Figure S3. Monocyte heterogeneity in the inflamed epidermis, related to Figure 2. A.** Heatmap showing scaled expression of differentially expressed genes between monocyte sub-clusters. **B.** Representative FACS gating strategy for GMP, MDP and monocytes isolation (red) from murine bone marrow for subsequent culture.

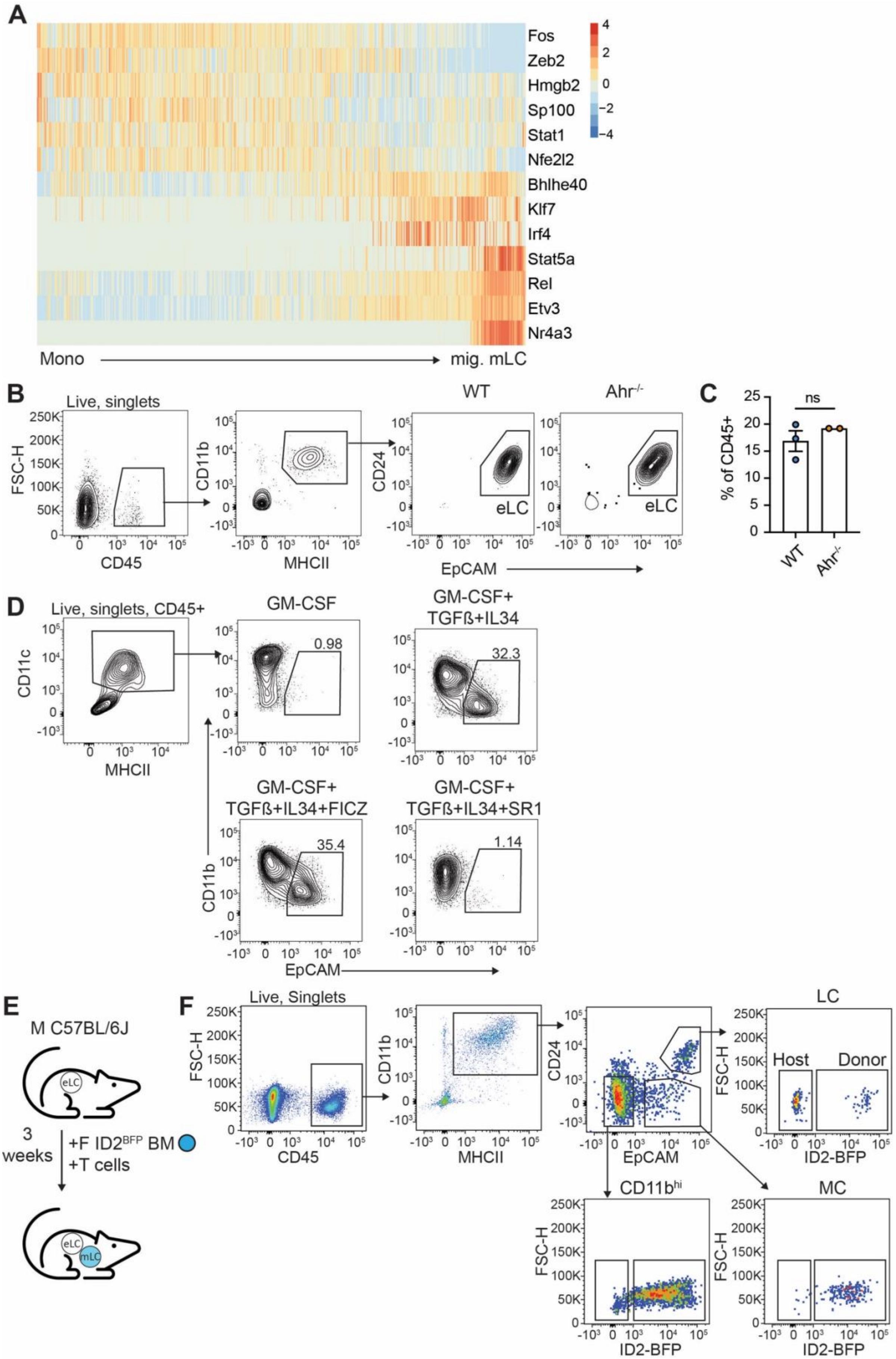

A
Fos
Zeb2
Hmgb2
Sp100
Stat1
Nfe2l2
Bhlhe40
Klf7
Irf4
Stat5a
Rel
Etv3
Nr4a3
Mono
mig. mLC
B
Live, singlets
FSC-H
CD45
CD11b
MHCII
WT
Ahr-/-
CD24
EpCAM
eLC
C
% of CD45+
ns
WT
Ahr-/-
D
Live, singlets, CD45+
CD11c
MHCII
GM-CSF
0.98
GM-CSF+ TGFß+IL34
32.3
GM-CSF+ TGFß+IL34+FICZ
35.4
GM-CSF+ TGFß+IL34+SR1
1.14
CD11b
EpCAM
E
M C57BL/6J
eLC
3 weeks
+F ID2BFP BM
+T cells
eLC
mLC
F
Live, Singlets
FSC-H
CD45
CD11b
MHCII
CD24
EpCAM
LC
Host
Donor
ID2-BFP
CD11bhi
MC

**Figure S4. Ahr in mLC differentiation *in vitro* and *in vivo,* related to Figure 4. A.** Heatmap showing scaled gene expression of transcription factors that are differentially expressed along the differentiation trajectory (Pseudotime) from monocyte to mig. mLC. **B.** Gating strategy and representative flow plots showing LC from WT and $Ahr^{-/-}$ murine epidermis. **C.** Bar graph showing frequency of LC from WT and $Ahr^{-/-}$ epidermis. Data are represented as mean±SD (n=3 for WT, 2 for $Ahr^{-/-}$). Statistical significance was assessed using a Mann-Whitney test. **D.** Gating strategy and representative flow plots showing mLC-like cells generated under indicated conditions. **E.** Diagram of experimental set up to track *in vivo* mLC differentiation following GVHD. Irradiated male B6 mice receive female $ID2^{BFP}$ bone marrow, CD4 T cells with Matahari T cells and LC chimerism is assessed by flow cytometry 3 weeks later. **F.** Representative gating strategy used to track *in vivo* mLC differentiation using $ID2^{BFP}$ reporter bone marrow.

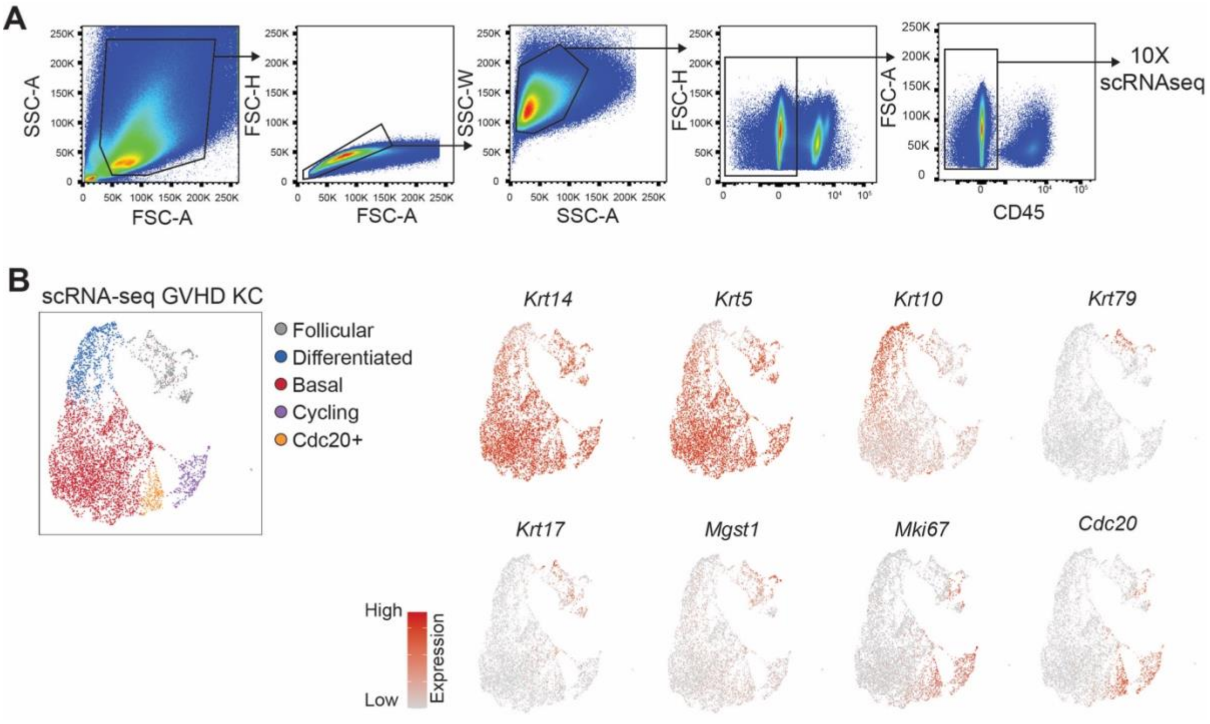


**Figure S5. scRNA-seq of keratinocytes from the GVHD epidermis, related to Figure 5. A.** Gating strategy for FACS isolation of keratinocytes from the GVHD epidermis for 10X scRNA-seq. **B.** UMAP visualisation and clustering of keratinocytes from GVHD epidermis analysed by scRNA-seq (left) and heatmap overlays showing expression of indicated genes across dataset (right). KC, keratinocytes.

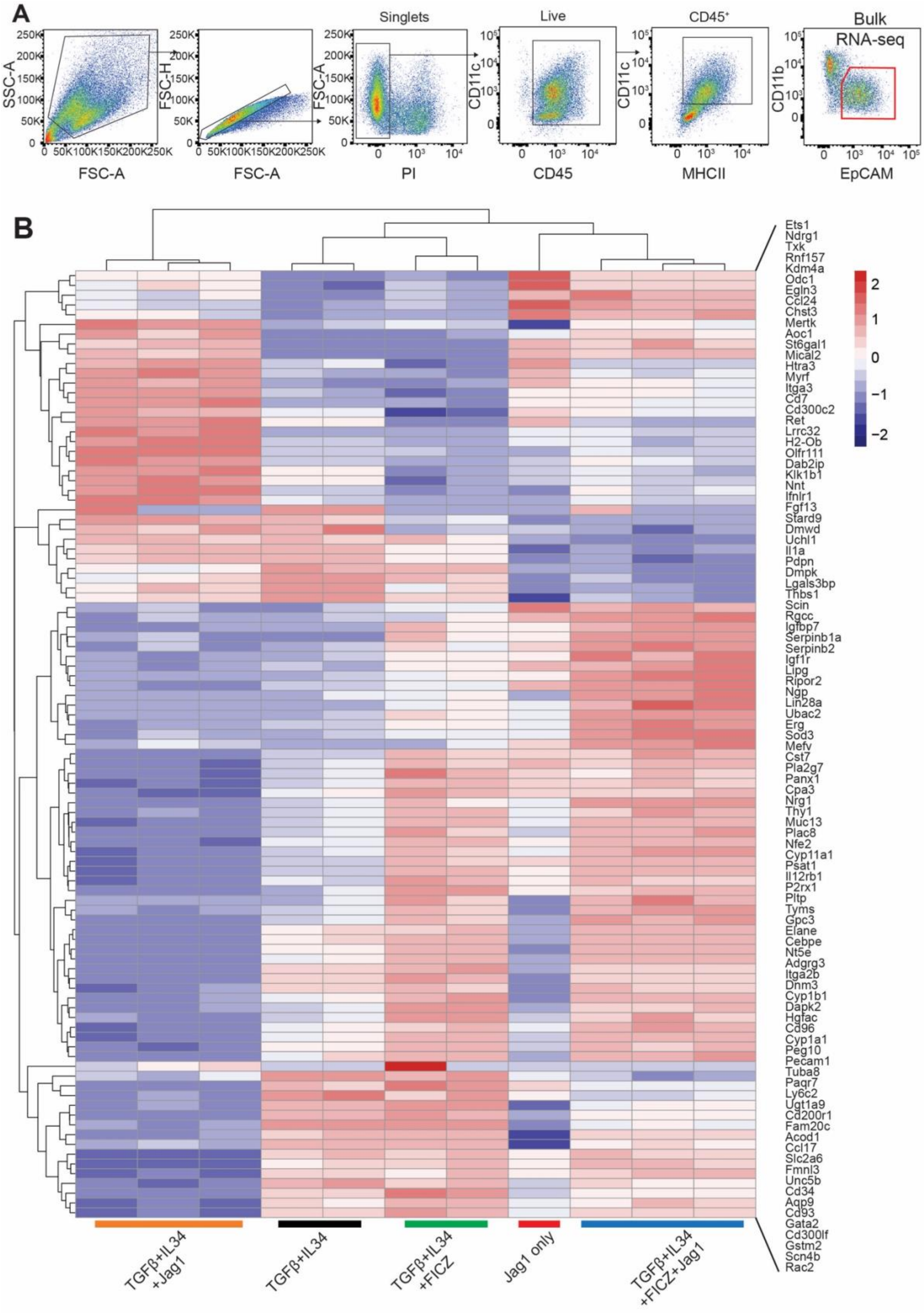

A
Singlets
Live
CD45⁺
Bulk RNA-seq
SSC-A
FSC-H
FSC-A
CD11c
CD11b
FSC-A
FSC-A
PI
CD45
MHCII
EpCAM
B
Ets1
Ndrg1
Txk
Rnf157
Kdm4a
Odc1
Egln3
Ccl24
Chst3
Mertk
Aoc1
St6gal1
Mical2
Htra3
Myrf
Itga3
Cd7
Cd300c2
Ret
Lrrc32
H2-Ob
Olfr111
Dab2ip
Klk1b1
Nnt
Ifnlr1
Fgf13
Stard9
Dmwd
Uchl1
Il1a
Pdpn
Dmpk
Lgals3bp
Thbs1
Scin
Rgcc
Igfbp7
Serpinb1a
Serpinb2
Igf1r
Lipg
Ripor2
Ngp
Lin28a
Ubac2
Erg
Sod3
Mefv
Cst7
Pla2g7
Panx1
Cpa3
Nrg1
Thy1
Muc13
Plac8
Nfe2
Cyp11a1
Psat1
Il12rb1
P2rx1
Pltp
Tyms
Gpc3
Elane
Cebpe
Nt5e
Adgrg3
Itga2b
Dnm3
Cyp1b1
Dapk2
Hgfac
Cd96
Cyp1a1
Peg10
Pecam1
Tuba8
Paqr7
Ly6c2
Ugt1a9
Cd200r1
Fam20c
Acod1
Ccl17
Slc2a6
Fmnl3
Unc5b
Cd34
Aqp9
Cd93
Gata2
Cd300lf
Gstm2
Scn4b
Rac2
2
1
0
−1
−2
TGFβ+IL34 +Jag1
TGFβ+IL34
TGFβ+IL34 +FICZ
Jag1 only
TGFβ+IL34 +FICZ+Jag1

**Figure 6. Bulk RNA-seq of in vitro mLCs, related to figure 6. A.** Representative gating strategy for FACS isolation of $CD11b^{int}EpCAM^{+}$ cells generated under conditions indicated in Figure 6C for bulk RNA-seq. **B.** Heatmap showing scaled expression of differentially expressed genes between the different indicated conditions.

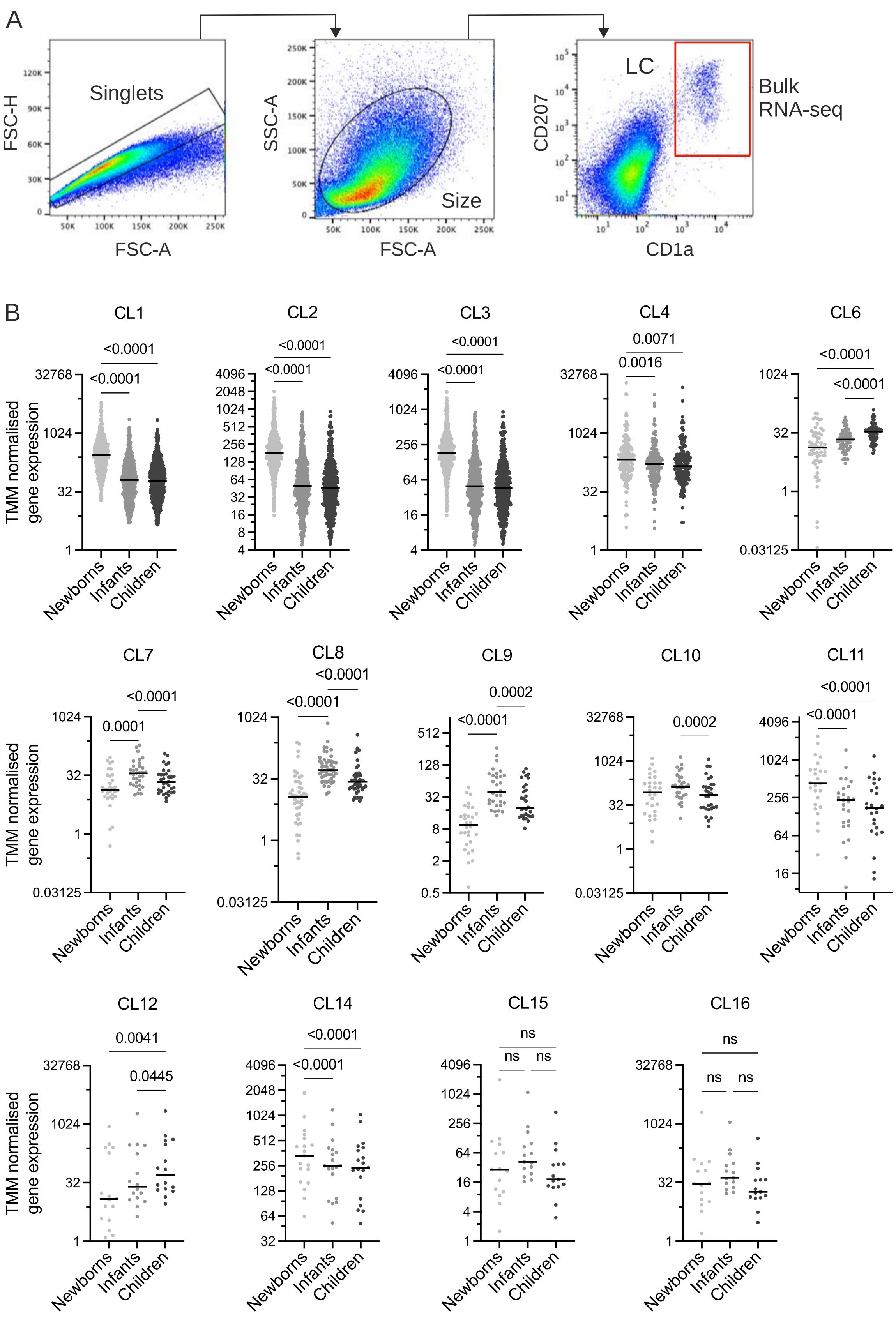
A
Singlets
FSC-H
FSC-A
SSC-A
Size
LC
CD207
CD1a
Bulk RNA-seq
B
CL1
CL2
CL3
CL4
CL6
CL7
CL8
CL9
CL10
CL11
CL12
CL14
CL15
CL16
TMM normalised gene expression
Newborns
Infants
Children
<0.0001
0.0071
0.0016
0.0001
0.0002
0.0041
0.0445
ns

**Figure 7. Bulk RNA-seq of human LCs from newborns, infants and children, related to Figure 7. A.** Gating strategy for cell sorting of human epidermis samples for LC bulk RNA-seq. **B**. Average TMM normalised gene expression levels in graphia clusters across newborns, infant and children.

**Supplementary Table 3.** List of murine antibodies used in this study.

| Antibody | Source | Identifier |
|---|---|---|
| Anti-mouse CD3e APC (145-2C11) | Biolegend | 100312 |
| Anti-mouse CD4 APC/Cy7 (GK1.5) | Biolegend | 100414 |
| Anti-mouse CD4 PE (GK1.5) | eBiosciences | 12-0041-82 |
| Anti-mouse CD4 PE/Cy7 (GK1.5) | Biolegend | 100422 |
| Anti-mouse CD8a v450 (53-6.7) | BD Biosciences | 560469 |
| Anti-mouse CD8a PE (53-6.7) | BD Biosciences | 12-0081-83 |
| Anti-mouse CD11b e450 (M1/70) | eBiosciences | 48-0112-82 |
| Anti-mouse CD11b APC (M1/70) | eBiosciences | 17-0112-83 |
| Anti-mouse CD11b PE (M1/70) | eBiosciences | 12-0112-82 |
| Anti-mouse CD11c FITC (HL3) | BD Biosciences | 553801 |
| Anti-mouse CD11c APC (HL3) | BD Biosciences | 550261 |
| Anti-mouse CD16/32 PE-Cy7 (93) | Biolegend | 101317 |
| Anti-mouse CD19 APC (1D3) | eBiosciences | 17-0193-82 |
| Anti-mouse CD24 FITC (M1/69) | Biolegend | 101805 |
| Anti-mouse CD24 BV650 (M1/69) | BD Biosciences | 563545 |
| Anti-mouse CD34 PE/Cy7 (SA376A4) | Biolegend | 152217 |
| Anti-mouse CD34 BV421 (SA376A4) | Biolegend | 152207 |
| Anti-mouse CD45 (30-F11) | Biolegend | 103155 |
| Anti-mouse CD45.1 BV605 (A20) | BD Biosciences | 563010 |
| Anti-mouse CD45.1 BV650 (A20) | Biolegend | 110736 |
| Anti-mouse CD45.2 PerCP/Cy5.5 (104) | BD Biosciences | 552950 |
| Anti-mouse CD45.2 APC/Cy7 (104) | Biolegend | 109824 |
| Anti-mouse CD49b APC (DX5) | Biolegend | 108910 |
| Anti-mouse CD64 BV786 (X54-5/7.1) | BD Biosciences | 741024 |
| Anti-mouse CD115 BV605 (AFS98) | Biolegend | 135517 |
| Anti-mouse CD117 BV510 (2B8) | Biolegend | 105839 |
| Anti-mouse CD135 PE (A2F10.1) | Biolegend | 135306 |
| Anti-mouse CD206 BV711 (CO68C2) | Biolegend | 141727 |
| Anti-mouse CD207 PE/Dazzle 594 (4C7) | Biolegend | 144211 |
| Anti-mouse B220 BV786 (RA3-6B2) | BD Biosciences | 563894 |
| Anti-mouse B220 APC (RA3-6B2) | Biolegend | 103211 |
| Anti-mouse EpCAM APC (G8.8) | eBiosciences | 17-5791-82 |
| Anti-mouse EpCAM PerCP/e710 (G8.8) | eBiosciences | 46-5791-82 |
| Anti-mouse Jagged-1 PE (HMJ1-29) | Biolegend | 130907 |
| Anti-mouse Ly6C PE-Cy7 (AL-21) | BD Biosciences | 560593 |
| Anti-mouse Ly6C FITC (AL-21) | BD Biosciences | 553104 |
| Anti-mouse Ly6G APC/Cy7 (1A8) | Biolegend | 127624 |
| Anti-mouse Ly6G APC (1A8) | Biolegend | 127613 |
| Anti-mouse MHCII BV510 (M5/114.15.2) | Biolegend | 107636 |

| | | |
|---|---|---|
| Anti-mouse MHCII APC/e780 (M5/114.15.2) | eBiosciences | 47-5321-82 |
| Anti-mouse MHCII A700 (M5/114.15.2) | Biolegend | 107621 |
| Anti-mouse Sca-1 BV711 (D7) | Biolegend | 108131 |
| Anti-mouse TER-119 APC (TER-119) | Biolegend | 116212 |
| Anti-mouse Vß8.3 TCR FITC (1B3.3) | BD Biosciences | 553663 |
| Anti-mouse Vß8.3 TCR PE (1B3.3) | BD Biosciences | 553664 |
